# Reconstruction of Par polarity in apolar cells reveals a dynamic process of cortical polarization

**DOI:** 10.1101/523589

**Authors:** Kalyn Kono, Shigeki Yoshiura, Ikumi Fujita, Yasushi Okada, Atsunori Shitamukai, Tatsuo Shibata, Fumio Matsuzaki

**Author notes:** equally contributed to this study (Shigeki Yoshiura, Ikumi Fujita).

## Abstract

Cellular polarization is fundamental for various biological processes. The Par network system is conserved for cellular polarization. Its core complex consists of Par3, Par6, and aPKC. However, the dynamic processes that occur during polarization are not well understood. Here, we artificially reconstructed Par-dependent polarity using non-polarized *Drosophila* S2 cells expressing all three components endogenously in the cytoplasm. The results indicated that elevated Par3 expression induces cortical localization of the Par-complex at the interphase. Its asymmetric distribution goes through three steps: emergence of cortical dots, development of island-like structures with dynamic amorphous shapes, repeating fusion and fission, and polarized clustering of the islands. Our findings also showed that these islands contain a meshwork of unit-like segments. Par-complex patches resembling Par-islands exist in *Drosophila* mitotic neuroblasts. Thus, this reconstruction system provides an experimental paradigm to study features of the assembly process and structure of Par-dependent cell-autonomous polarity.

## INTRODUCTION

Polarization is a fundamental cellular property that plays a vital role in various biological processes in multi-cellular as well as single cell organisms. Par-complex system is a conserved mechanism that regulates cell polarization(Kemphues *et al*, 1988; Suzuki & Ohno, 2006; Johnston, 2018). The core Par-complex consists of Par6, Par3, and typical protein kinase C (aPKC) (Kemphues *et al*, 1988; Tabuse *et al*, 1998). Domain structures of these components and their interactions have been extensively studied(Lang & Munro, 2017). Par3 exhibits membrane binding affinity through its C-terminal domain and the ability to self-oligomerize via its N-terminal CR1 domain, which is essential for its localization and function(Benton, 2003; Mizuno *et al*, 2003; Krahn *et al*, 2010; Harris, 2017) Structural studies have revealed that the CR1 domain forms helical polymers of 10 nm diameter(Zhang *et al*, 2013). Par6 and aPKC, which form a stable subcomplex, interact with the CR3 and PDZ domains of Par3(Izumi *et al*, 1998; Renschler *et al*, 2018). Phosphorylation of this domain by aPKC inhibits this interaction(Morais-de-Sá *et al*, 2010; Soriano *et al*, 2016). Thus, Par-complex assembly is a dynamic process. CDC42 binds to the aPKC-Par6 subcomplex and anchors it to the cell membrane as a diffusible cortical form(Joberty *et al*, 2000; Aceto *et al*, 2006; Rodriguez *et al*, 2017; Wang *et al*, 2017) On the other hand, Lgl and/or Par1 kinase act as inhibitory factors against aPKC(Guo & Kemphues, 1995; Betschinger *et al*, 2003; Yamanaka *et al*, 2003; Plant *et al*, 2003; Hurov *et al*, 2004), and distribute complementarily to the core Par complex. Interplay between these components results in cytocortical asymmetry(Doerflinger *et al*, 2006; Sailer *et al*, 2015).

Cell polarization involving the Par-complex *in situ* is linked to various other processes. The Par-complex creates epithelial cell polarity during interphase at the subapical domain (including tight junctions) that is tightly associated with adherens junctions, where Par3 primarily localizes(Rodriguez-Boulan & Macara, 2014; Suzuki & Ohno, 2006). On the other hand, cell polarization is coupled with mitosis during asymmetric divisions, and autonomously induced or triggered by an external cue, depending on the cell type(Yamashita *et al*, 2010). Because of such association between Par-dependent polarization and other processes, the Par-complex exhibits different behavioral characteristics in an individual context, making it difficult to determine general features of the dynamic process taking place during cell polarization by the Par-complex. We attempted to address this problem by establishing an artificial polarization system induced by the Par-complex(Baas *et al*, 2004; Johnston *et al*, 2009). We used *Drosophila* Schneider cells (S2 cells) of mesodermal origin as host cells(Schneider, 1972). They are neither polarized nor adhere to the substratum and between cells. The 3 core components of the Par-complex are endogenously expressed in S2 cells but are distributed in the cytoplasm throughout the cell cycle. Thus, S2 cells appear to be an ideal system for cell polarity induction.

## RESULTS

### S2 cells polarize by an elevated expression of Par3

First, we tested the effect of overexpressing each core component of the Par-complex in S2 cells, which distribute these components evenly throughout the cytoplasm and divide symmetrically (Fig. 1**a**). We found that all core components of the Par-complex cortically co-localized in an asymmetric manner when Par3 was over expressed, but did not cortically localized, when Par6 or aPKC was overexpressed (Figs. 1**b**, **c**, and data not shown). We overexpressed myc-Par3 (or Par3-mKate2) via the *actin*-promoter (*act5c*)-driven *Gal4-UAS* system (*Act-Gal4>UAS*) by transfection (see Methods), with or without *actin*-promoter-*Par6-GFP* (*pAct-Par6-GFP*) as a live marker, which was uniformly distributed in the cytoplasm in the absence of Par3 overexpression (Fig. 1**b**). Among transfected cells that exhibited cortical Par complex distribution, a fraction exhibited an asymmetrically localized Par-complex (Fig. 1**c**, Supplementary Fig. 1**b**), while the rest of the cells localized uniformly to the cortex (see below). Asymmetric distribution of the Par-complex induced by Par3 overexpression required endogenous aPKC and Par6 (Fig. 1**d**). The expression level of exogenous Par3 was also important for S2 cell polarization. Cortical polarization was not observed (Fig. 1**b**) when Par3 expression, directly driven by the *actin* promoter, was approximately 1/40 of that of the *Act-Gal4>UAS* system (Supplementary Fig. 1**a**).

**Figure 1.**
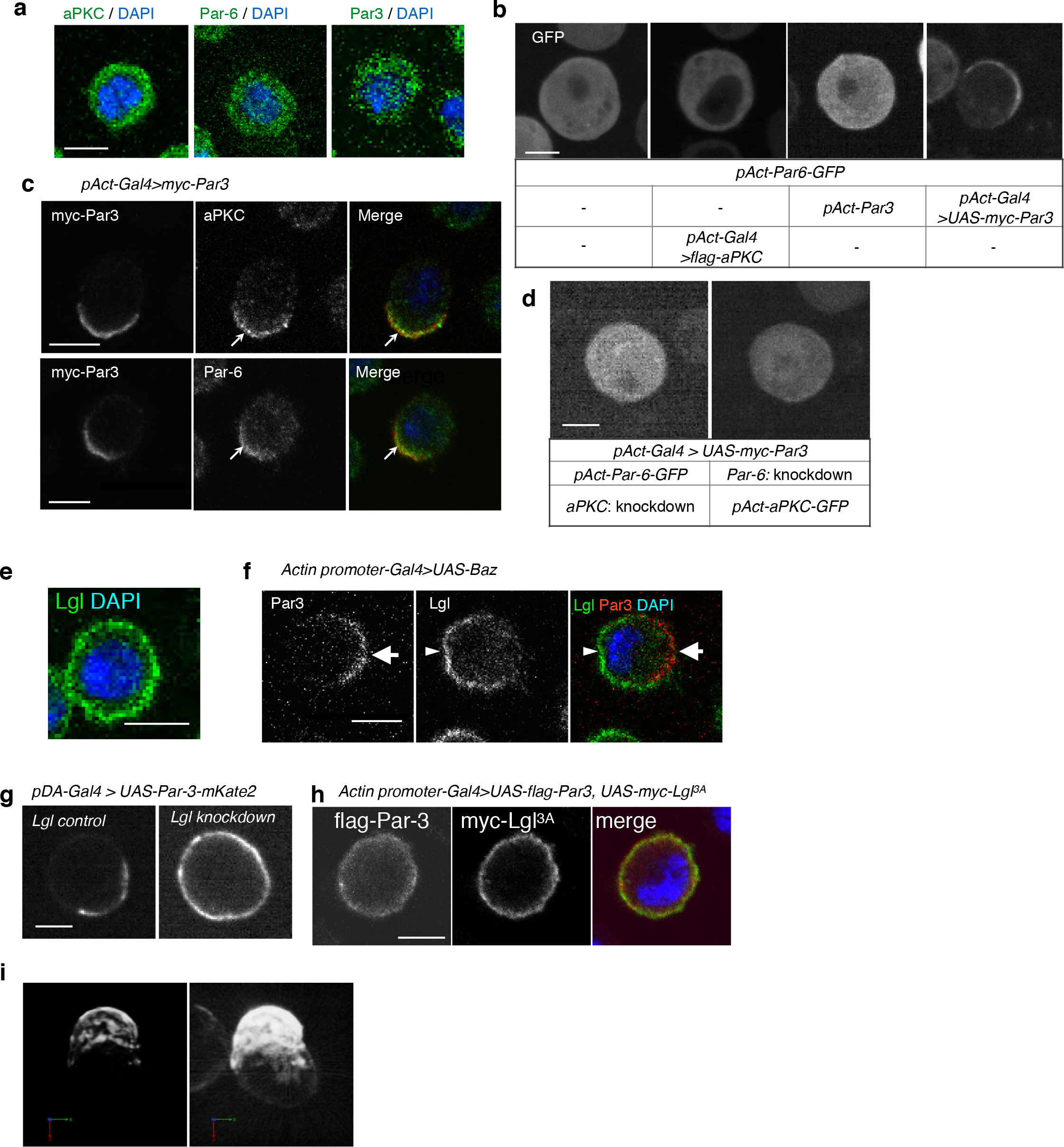
S2 cells polarize due to elevated Par3 expression. **a.** Immunostaining of endogenous aPKC, Par6, and Par3 in S2 cells two days following transfection of the empty vector. Blue indicates DAPI staining. Images in a-i were at the equatorial plane of cells. Scale bar, 5 μm in all panels in this figure. **b.** Live-imaging of Par6-GFP in S2 cells (top), two days following transfection of a combination of expression plasmids as described in the table (bottom). **c.** Localization of endogenous aPKC and Par6 in cells overexpressing myc-Par3, stained with anti-myc-tag and anti-aPKC or anti-Par6 antibodies, and with DAPI, two days after transfection. Arrows indicate co-localized Par components. **d.** Live-imaging of Par6-GFP (left) or aPKC-GFP (right) in Par3-overexpressing cells containing aPKC or Par6 RNAi knockdown, respectively, at two days post-transfection. **e.** Endogenous expression of Lgl in S2 cells stained with anti-Lgl and DAPI at two days post-transfection of the empty vector. **f.** Par3 and endogenous Lgl localize complementarily in 71% of cells (n=24) where overexpressed Par3 was asymmetrically localized. Arrow, Par3 crescent. Arrowhead, Lgl. **g.** Live-imaging of myc-Par3-mKates without (left) or with (right) Lgl knockdown by RNAi at two days post-transfection. **h.** S2 cells over-expressing flag-Par3 and myc-Lgl3A, stained with anti-flag-tag, anti-myc-tag and DAPI. Lgl3A was cortically uniform in contrast to cytoplasmic Par3 distribution. **i.** 3-D reconstructed image of a cell overexpressing myc-Par3-mKate2 (left). In the right side image, brightness and contrast were adjusted to visualize the outline of the same cell. The movie of a different cell is shown as Supplementary movie 1.

We next examined the effect of endogenous Lgl that was largely localized uniformly along the cortex with a cytoplasmic distribution in S2 cells, prior to Par3 overexpression (Fig. 1**e**). When Par3 was distributed asymmetrically along the cortex, Lgl and Par3 distributed in a complementary manner (Fig. 1**f**). Knockdown of *lgl* via RNAi and the expression of Lgl3A, which aPKC is not able to phosphorylate, showed that Lgl and its phosphorylation by aPKC are required for asymmetric Par-complex localization in S2 cells (Fig. 1**g**,**h**).

To evaluate the degree of polarization of transfected cells, we introduced the asymmetric index (ASI), a measure of the polarized Par-complex distribution, which, according to Derivery et al.(Derivery *et al*, 2015), indicates the degree of polarization of a fluorescent marker distributed along the equatorial cortex (Supplementary Fig. 1**b**). ASI distribution was compared with that of membrane-bound GFP (memGFP), which is essentially non-polarized (the control). The ASI value of memGFP ranged from 0 to 0.35 due to fluctuation. Par3 cortical distribution was categorized into 2 groups in comparison with memGFP. Cells with an ASI in the same range as that of memGFP were regarded as non-polarized. Those cells showing an ASI larger than that of the mem-GFP were interpreted as polarized. Among transfected cells showing cortical distribution of the Par complex, 38% were polarized, while 62% were non-polarized (Supplementary Fig. 1**c)**.

Furthermore, we examined the 3-dimensional (3D) distribution of the Par complex in S2 cells by reconstructing serial images at steady state two days following transfection. Interestingly, the region where the Par-complex accumulated was not uniform but consisted of multiple large aggregates (Fig. 1**i**). These large aggregates were termed “Par-islands”. These islands dynamically changed their arrangement on the surface of the S2 cells (Supplementary movie 1).

### Temporal patterns of Par-complex polarization

To investigate temporal patterns of polarized distribution of the Par-complex, we induced Par3 expression via the *Metallothionein* promoter, which is activated in the presence of CuSO_4_. We transfected S2 cells with *pMT-myc-Par3-mKate2, pAct-Par6-GFP* and *pAct-aPKC*. Because expression of Par6-GFP and aPKC reached steady levels two days following transfection (Supplementary Fig. 2**a**), induction of Par3-mKate2 was initiated at this time. This indicated that the Par-complex first emerged in dot form (designated Par-dot) in the cell cortex during the initial phase of myc-Par3-mKate2 elevation, 2 -3.5 h following induction (Fig. 2**a-c**, Supplementary movie 2). Interestingly, Par-dots emerged in a restricted region of the cell cortex (Fig. 2**c**).

**Figure 2.**
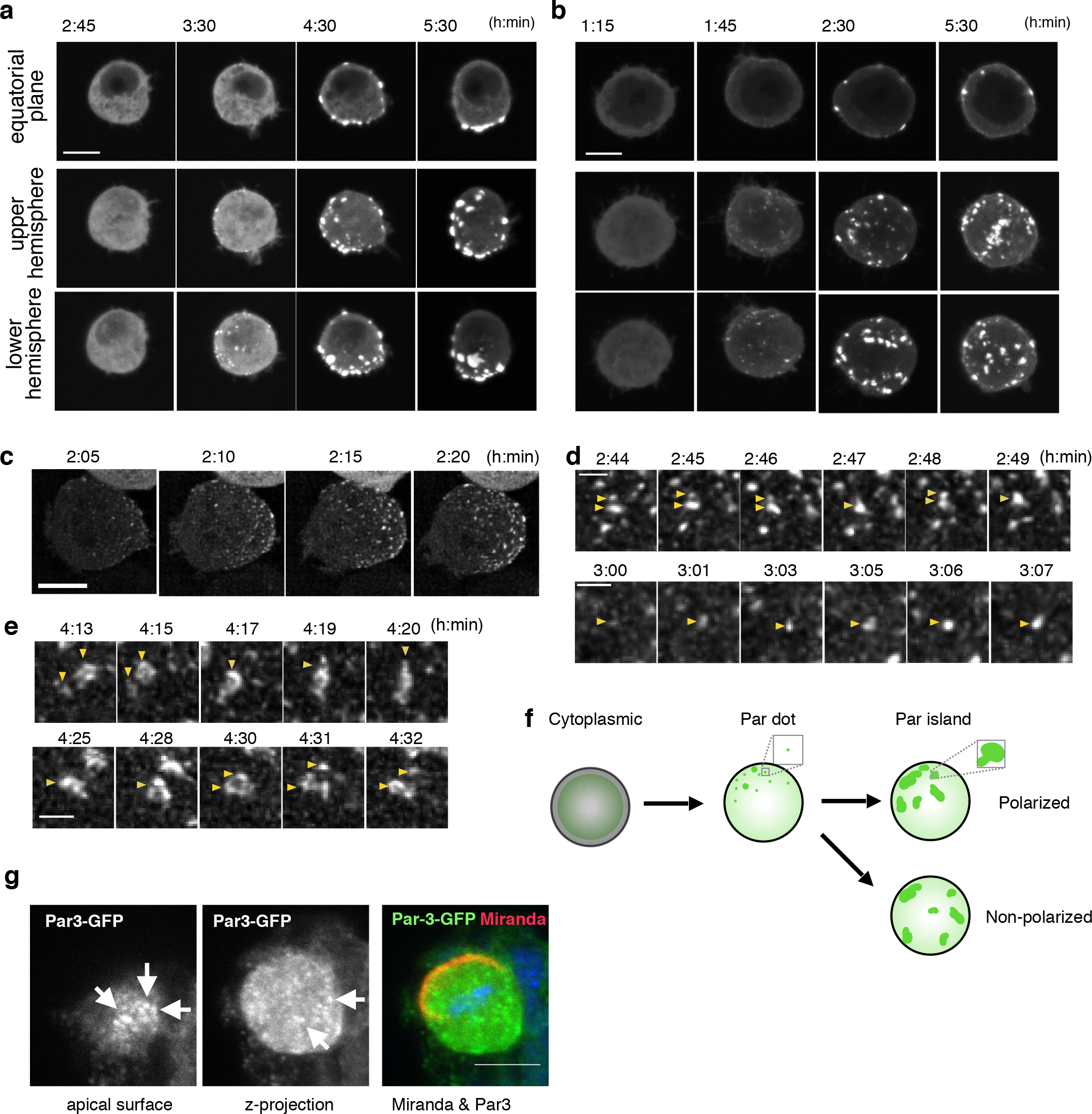
Temporal pattern of Par complex clustering. **a, b**. Time-lapse images of S2 cells inducing Par3-mKate2 expression via the *Metallothionein* promoter, leading to polarized (**a**) or non-polarized (**b**) Par3 distribution. Time 0 (h: min) was at the time of induction by CuSO_4_ addition in these and subsequent panels. The top row shows images at the equatorial plane. The middle and bottom rows show the max intensity projection images of the upper and lower hemispheres of the cell, respectively. Scale bar, 5 μm. **c.** Time-lapse images of Par6-GFP showing the emergence and development of Par-dots. The images are 6 μm max intensity projection covering the entire cell. Scale bar, 5 μm. **d.** Time-lapse imaging of Par6-GFP showing the fusion and fission of Par-dots (arrowheads in the upper panel), and the growth of a Par-dot (arrowheads in the lower panel). In **d** and **e**, scale bar, 1 μm. **e.** Time-lapse image of Par-islands visualized by Par6-GFP. Arrowheads indicate dynamic shape changes, fusion (arrowheads, upper panel) and the dissociation of Par-islands (arrowheads, lower panel). **f.** Schematic presentation of S2 cell polarization process from Par-dot formation to clustering of Par-islands. **g.** Localization of the Par3-GFP in a mitotic neuroblast of a *Drosophila* brain expressing *Par3-GFP*. A brain taken from a 3^rd^ instar larvae and stained for GFP and Miranda(Ikeshima-Kataoka *et al*, 1997). A single plane near the apical surface (left), A max-intensity projected image of the whole cell (center) and merged (right) image are shown. Par-island like structures are observed (arrows). Scale bar, 5 μm.

Par-dots continued to grow in size via self-expansion, repeated fusion and less frequent fission (Fig. 2**d**). These dots then developed into “islands” of various sizes and shapes 2.5-6 h following induction (Figs. 2**a,b**), as observed in the *UAS-Gal4* system (Fig. 1**i**). During this period, the distribution of Par-islands occurred via 2 separate processes, polar and non-polar clustering, which corresponded with temporal changes in ASI values (Figs. 2**a,b;** Supplementary Fig. 2**e**). However, there was no significant difference between polarized cells and non-polar cells in either the time course of Par3-mKate2 expression levels or the Par6-GFP/Par3-mKate2 ratio (Supplementary Figs. 2**b-d**). While the average steady state amount of Par3-mKate2, driven by the *Metallothionein* promotor was approximately 1/18-fold of that driven by the *Act-Gal4>UAS* system (Supplementary Fig. 2**f**), the appearance of the islands was similar between the two expression systems (Figs. 1**i** and 2**a**).

Island structures of the Par-complex were also observed in the apical domain of *Drosophila* mitotic neuroblasts (Fig. 2**g**), suggesting that formation of Par-islands may be a common process during the polarized cortical distribution of the Par-complex irrespective of cell cycle phases.

### Dynamics at the steady state

Once the Par3-mKate2 expression level steadied about 8 h following induction, approximately 30% of cells with Par6-GFP localized cortically demonstrated polarized distribution of Par6-GFP, resulting in a crescent in the equatorial plane (ASI > 0.35), while the rest of cells showed a non-polarized cortical distribution (ASI≦0.35); (Fig. 3**a**). At steady state, Par-islands dynamically change their mutual positions (Fig. 3**b**, Supplementary movie 3). However, their asymmetric clustering and non-polar distribution were largely maintained for at least several hours, once cells reached steady state (Supplementary Fig. 2**g**), suggesting that both polar and non-polar clustering of Par-islands was fairly stable. In both states of Par-island clustering, the distribution of Par-islands and Lgl in S2 cells were mutually exclusive (Figs. 3**c,d**, Supplementary movies 4, 5), suggesting that Lgl plays a role in the stability of the 2 states. Interestingly, Par-islands were never unified into one large island regardless of their dynamic movements as well as fusion and fission (Fig. 3**b**).

**Figure 3.**
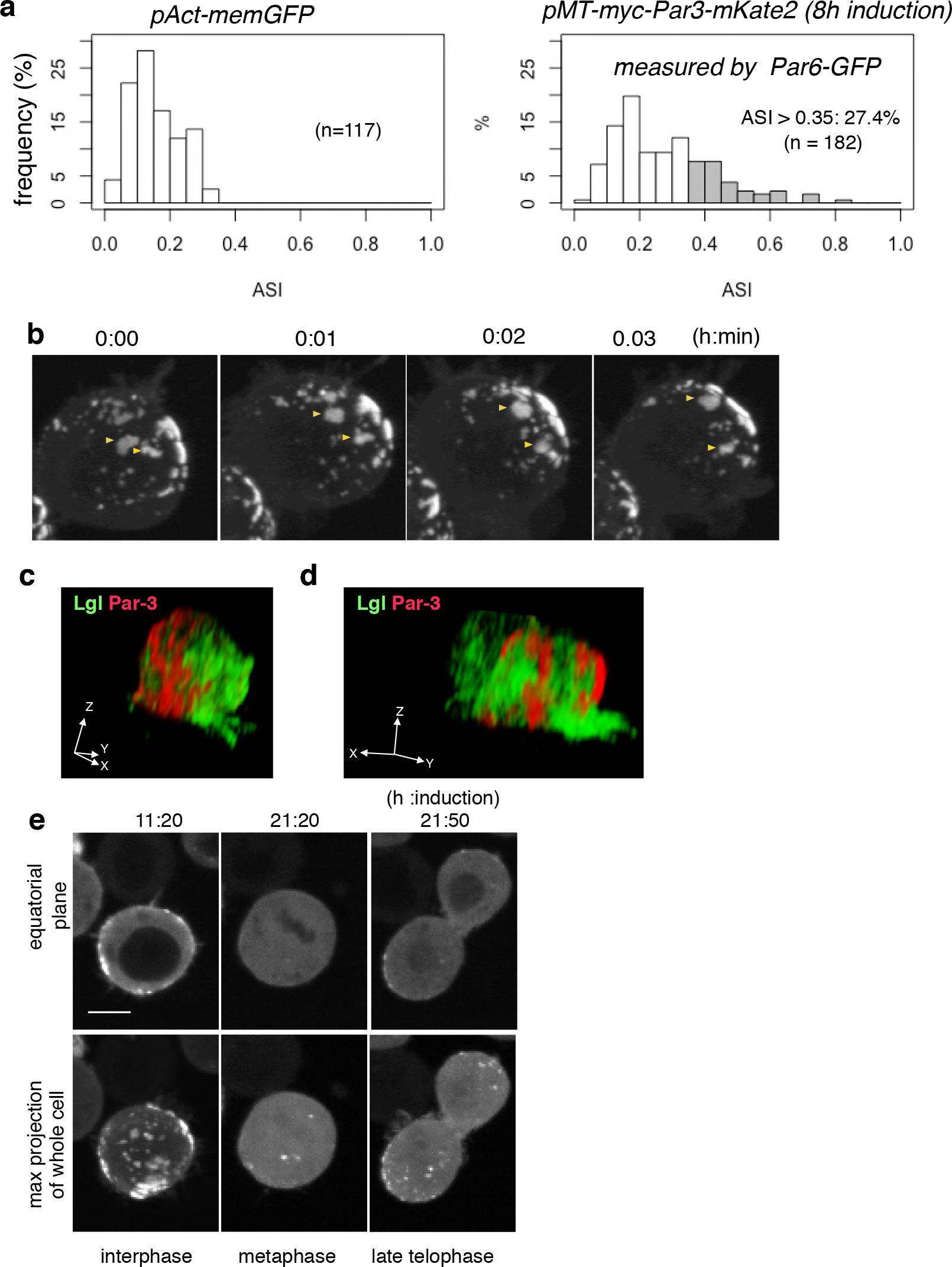
Steady state dynamics of polarized Par complex. **a.** The distribution of ASI among cells with memGFP (left) driven by the *Act5C* promoter and Par-3-mKate2 (right) induced MT promoter. ASI was measured for the equatorial plane of cells 8 h after CuSO_4_ addition. The mean ASI value was 0.16±0.07 (s.d.) for cells expressing memGFP (n = 117 cells), and 0.27±0.15 (s.d.) for cells localizing Par3-mKate2 along the cell cortex (n=182 cells). Cells showing ASI in the range outside the ASI distribution for memGFP expressing cells (ASI>0.35) were 27.4 % of the cells with cortical Par3-mKate2. Mean ASI value for those cells was 0.43±0.12 (s.d.). In all figures and the main text, s.d. is shown following the mean value. **b.** Time-lapse imaging of Par-islands at the steady state, taken 8 h after CuSO_4_ addition and onwards. **c** and **d**. 3D images of the distribution of myc-Par3-mKate2 and endogenous Lgl in cells showing polarized (**c**) and non-polarized (**d**) Par3 localization. The distribution of Lgl and myc-Par3-mKate2 is essentially non-overlapped in both cases. See Supplementary movie 4 for the 3D-rotation movies. **e.** Time-lapse imaging of Par3 distribution during mitosis. Time indicates h:min after CuSO_4_ addition. Images of equatorial plane (upper panels), and the max projection of the whole cell (lower panels) are shown. Scale bar, 5 μm.

Upon mitosis, cortical Par-islands disassembled and disappeared (Fig. 3**e**). However, just prior to cell cleavage, small Par-dots reappeared preferentially in the region opposite the cleavage site (n=13/15). Consistently, the position of the centrosome, which is normally located on the far side from the cleavage site, coincided weakly with the region of Par-dot reappearance (Supplementary Figs. 2**h-j**).

### Structural analysis of the assembly state of the Par-complex

The island form of the Par-complex is a unique structure with a slightly convex shape (Figs. 2**a,b**, 3**b**, Supplementary movie 3). To better understand the organization of Par-islands, we investigated their structure at the super-resolution level. We performed super-resolution radial fluctuation (SRRF) analysis(Gustafsson *et al*, 2016), using confocal images of fixed samples, double-stained by Par3-mKate2 and Par6-GFP. This analysis revealed that both Par3-mKate2 and Par6-GFP exhibited polygonal shaped islands of various types (Figs. 4**a,b**, Supplementary Figs. 3**a, b**). In order to determine whether there was regularity in these structures, we measured contour lengths of the Par3-mKate2-stained meshwork, including separate rods and polygons. This measurement yielded distribution of lengths with multiple peaks in the density plot. We then searched the regularity of these multiple peaks. We found that it was well fitted with a combination of seven Gaussian curves, which exhibited a peak interval of 0.38±0.06 μm (mean ± s.d., hereafter); (Figs. 4**c, d**). We also performed spectral analysis of the density plot, and obtained a single major frequency of 2.4 μm^−1^, which gives a peak interval of 0.42 μm (Fig. 4**e**). These two analyses thus give consistent results with each other.

**Figure 4.**
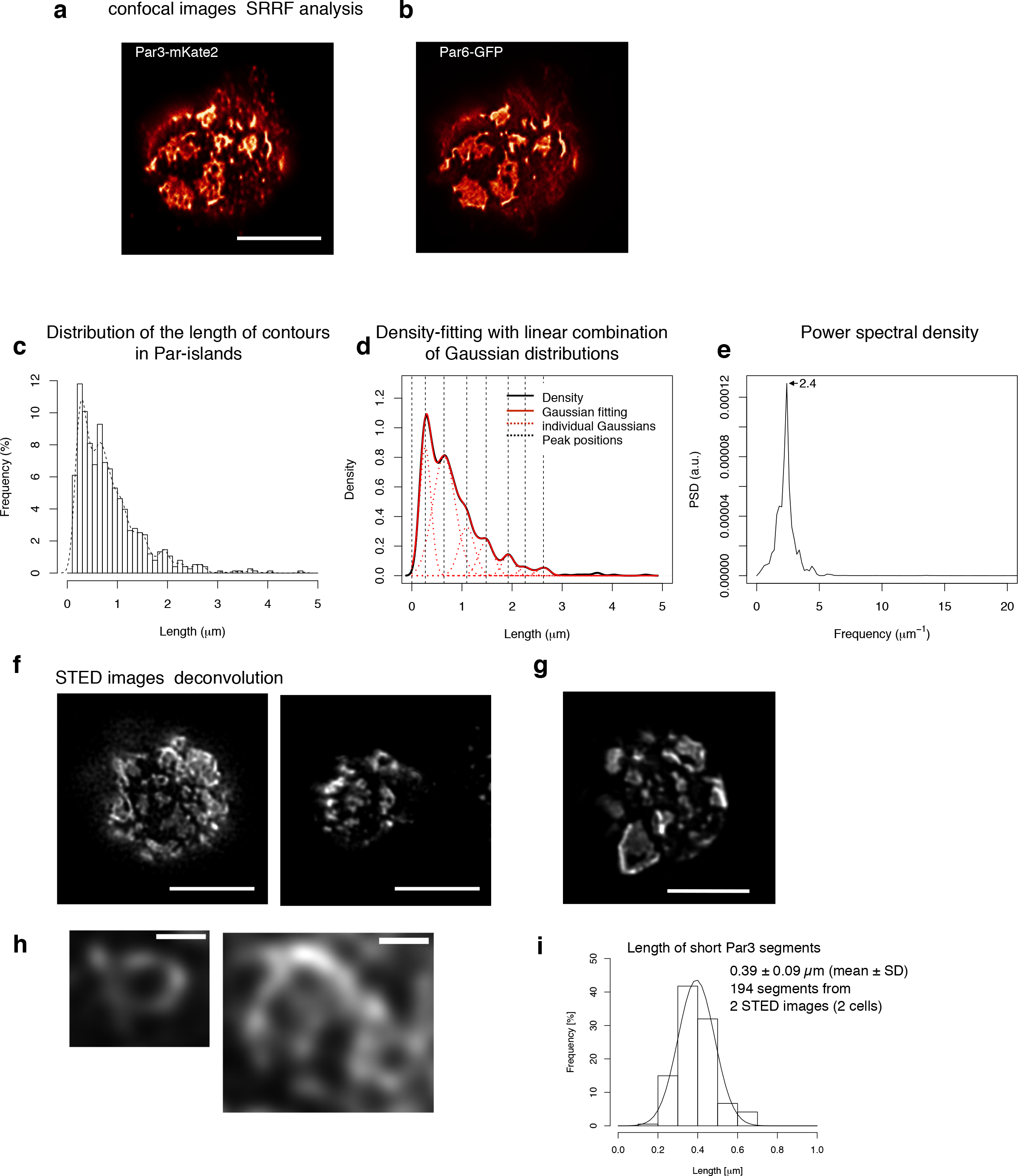
SRRF-processed confocal images and STED microscopy reveals a unit-like segment in Par-islands. **a** and **b**. SRRF-processed confocal images of cells expressing both Par3-mKate2 (**a**) and Par6-GFP (**b**). Scale bar, 10 μm. **c.** Histogram of the distribution of contour lengths along the meshwork in Par-islands visualized by Par3-mKate2. Cells were transfected with pMT-Par3-mKate2 and pAct-Par6-GFP, and treated with CuSO_4_ as described above. The contour lengths were measured following edge detection processing of SRRF-processed images (see Supplementary Fig. 3**a** and Methods for actual measurements). **d.** Gaussian fitting of the density plot (**c**). The density plot of the histogram (**c**) was fitted with 7 Gaussian curves via the least square method. Parameters of individual Gaussian curves are shown in Supplementary Fig. 3**b**. The averaged mean of individual Gaussian curves was 0.38±0.062 μm for 754 contours from 28 cells. **e.** Power spectral density for the second derivative of the contour distribution plot shown in (**c**). The major frequency was 2.4 μm^−1^. **f** and **g**. Deconvoluted super-resolution (STED) images of cells expressing myc-Par3-GFP (**e**) and Par6-GFP (**f**), stained for GFP. S2-cells were transfected with pMT-myc-Par3-GFP and pAct-Par6 (e) or with pMT-myc-Par3 and pAct-Par6-GFP (fb), followed by CuSO_4_ addition for induction two days following transfection. Cells were fixed for immune-staining for GFP at 8 h post-induction. Scale bar, 10 μm. **h.** Magnified views of the left cell in (**e**). Scale bar, 0.5 μm. **i.** Distribution of the length of individual segments constructing Par-islands visualized by Par3-GFP and Gaussian fitting. See Supplementary Fig. 3**c-f** for measurements. The mean value of the single segment lengths was 0.39±0.09 μm based on 194 segments from 2 STED images for 2 cells (**a**).

We also observed Par-islands of Par3-GFP and Par6-GFP separately via STED microscopy, and found similar meshwork structures in the deconvoluted STED images (Figs. 4**f-h**). The linear part of segments in the meshwork structures were measured (Supplementary Figs. 4**a-d**), and exhibited a length of 0.39±0.09 μm (Fig. 4**i**). Thus, these two methods essentially provided the same value for the segmental length. In addition, these segments had a fairly homogeneous diameter in STED microscopic images, where the mean half width of Par-segments was 0.22±0.03μm, (Supplementary Figs. 4**e, f**). These results raise the possibility that the Par-island meshwork contains a unit segment. Indeed, separate rod- or string-shape structures as well as open square-structures were often observed in the earlier phases of the Par-complex aggregation time course (Supplementary Fig. 4**g**, movie 2), supporting the notion that Par-islands are assembled from these elemental structures, generating regularity in the meshwork organization.

### Roles of Par components and the cytoskeleton in polarity formation

Because the elevation of Par3 expression induced cortical polarization in S2 cells, we investigated the role of functional domains of Par3 by observing phenotypes with Par6-GFP following the overexpression of mutant *Par3* forms via the *Metallothionein* promoter (Fig. 5**a**). First, we tested the role of the CR1 domain responsible for self-polymerization in the polarized Par-complex assembly(Benton, 2003; Mizuno *et al*, 2003). Overexpression of Par3 lacking CR1(Par3ΔCR1) in the presence of the endogenous Par3 compromised the cortical Par-complex assembly significantly. The Par-complex was broadly distributed at the cell cortex in the initial stages, when dots were very faintly visible. While denser parts similar to Par-islands formed later, they were mostly faint with ambiguous contours, compared to those formed by wild type Par3 expression (compare Figs. 2**a, b** and Fig. 5**b**), and the Par-island distribution was not eventually polarized (Fig. 5**e**). These results suggested that the CR1 domain was important for all processes during the development of macro-scale structures of the Par-complex. However, this phenotype was different from that of the non-transfected S2 cells, suggesting that a large amount of Par3ΔCR1 contributes to cortical Par-complex aggregation in the presence of endogenous Par3.

**Figure 5.**
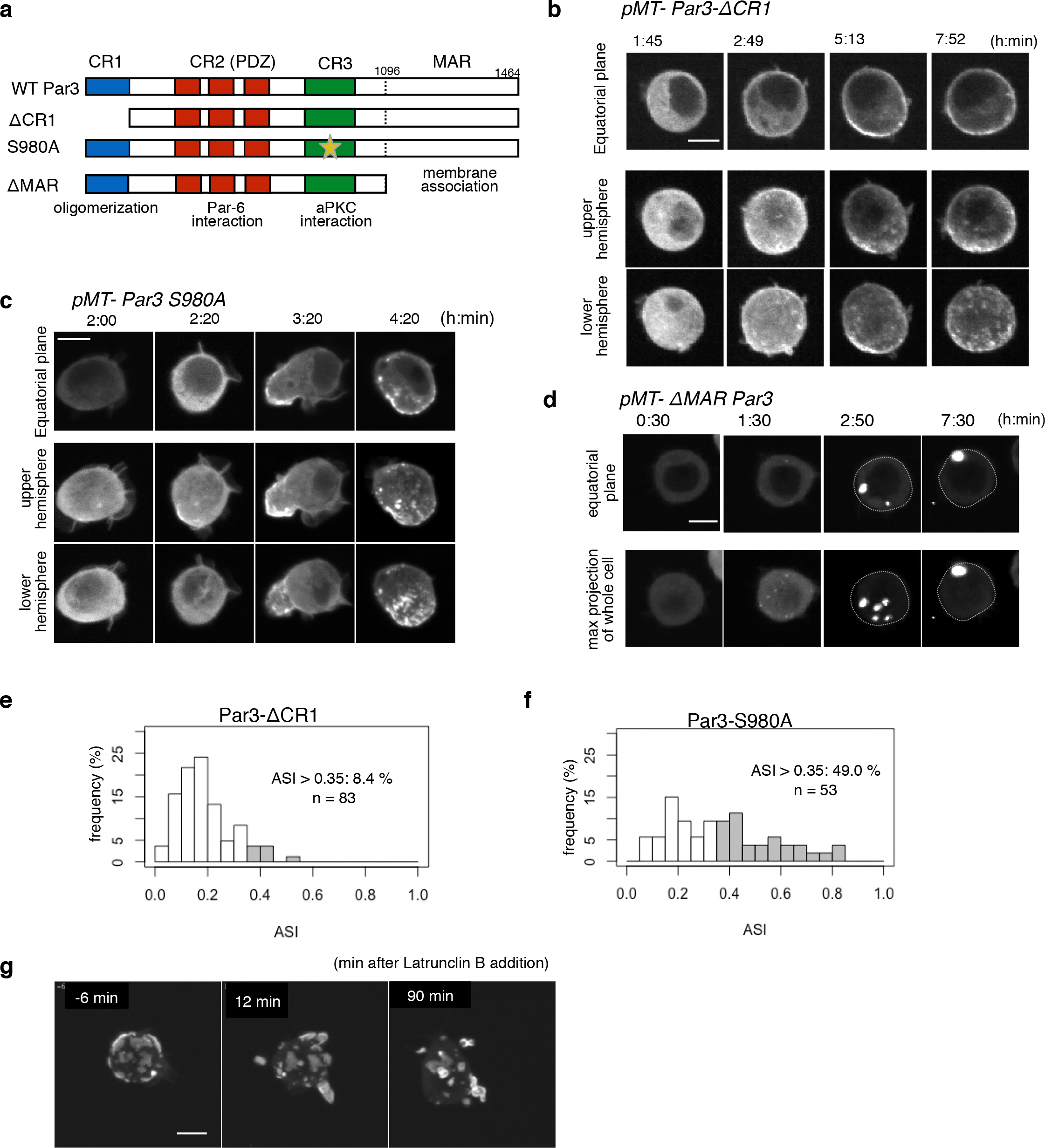
Role of functional domains of the core Par complex. **a.** Schematic representation of the functional domains of Par3 and their corresponding mutant constructs used in this study. **b-d**. Time-lapse imaging of the distribution of Par6-GFP in cells where Par3^ΔCR1^-mKate2 (**b**), Par3^S980A^-mKate2 (**c**) and Par3^ΔMAR^-mKate2 were induced by the *Metallothionein* promoter. Time is indicated in h:min after CuSO_4_ addition. In (**b**) and (**c**), images at the equatorial plane (top panels), and stacked images of the upper hemisphere (middle panels), and the lower hemisphere (bottom panels) are shown. In (**d**), the lower panels show maximum-projection images of the whole cell. Scale bar, 5 μm. **e** and **f**. The distribution of ASI is shown for cells that have induced Par3^ΔCR1^-mKate2 (**e**, the mean value = 0.19±0.10 for n = 83 cells), and Par3^S980A^-mKate2 (**f**, the mean value = 0.37±0.20 for n = 53 cells). The gray part of histograms indicates the fraction of cells having ASI in the range out of the ASI distribution for memGFP-expressing cells (ASI > 0.35, see Supplementary Fig. 1**c**, and Fig. 3**a**). Cells in this range are 8.4 % of cells with cortical Par3^ΔCR1^-mKate2 (mean ASI value = 0.41±0.06) and 49.0% for Par3^S980A^-mKate2 (mean ASI value = 0.54±0.14). In both cases, the polarized cell population (ASI >0.35) is significantly altered (p=0.0006 for Par-3^ΔCR1^-mKate2 and p=0.0088 for Par-3^S980A^-mKate2) compared with that of wild type Par3-mKate2 (Fig. 3a). Quantification was performed 8 h after CuSO_4_ addition onwards. In all images, CuSO_4_ was added at two days post-transfection. **g.** Time-lapse imaging of the effects of actin disruption on the Par-islands. At 8h after Par3-mKate2 induction, cells were treated with Latrunculin B. Par-islands rapidly became round and/or promoted membrane protrusion. Faint fluorescent islands face the bottom of the dish. See Supplementary movie 6. Scale bar, 5 μm.

We next examined the effect of aPKC-dependent phosphorylation at Serine 980 in the CR3 domain, which is necessary for dissociation of Par3 from aPKC (Fig. 5**a**)(Morais-de-Sá *et al*, 2010). Overexpressing the non-phosphorylatable form, Par3S980A, which tightly binds aPKC(Morais-de-Sá *et al*, 2010), increased the polarized cell population, where 49% of cells with cortical Par3 showed an ASI > 0.35, and a degree of polarization with a mean ASI value of 0.54±0.14 for polarized cells. This suggested strong enhancement of Par-complex clustering (Figs. 5**c, f**). Clustering of the Par-islands was so tight under this condition that the polarized region sometimes assumed a bowl-like shape, in which the island structure was hardly discernible. Subsequently, this dense aggregation gradually separated into small and nested islands. Dense packing of the Par complex containing Par3S980A suggested that the turnover of Par3-aPKC association and dissociation played a role in the normal clustering of Par-islands. This was similar to that of *Drosophila* epithelial cells, wherein Par3S980A colocalized with aPKC-Par6 in the apical domain with disorganized adhesion belts(Morais-de-Sá *et al*, 2010).

Next, we examined the effect of the membrane association region (MAR) of Par3 by overexpressing Par3ΔMAR(Krahn *et al*, 2010). The Par-complex no longer localized cortically, but formed several cytoplasmic aggregates, which coalesced into a single large sphere (Fig. 5**d**). Thus, the functional domains of Par3 and the interactions between these domains, together, play a role in the properly polarized distribution of the Par-complex in the S2 cell system.

Lastly, we examined the effects of the actin cytoskeleton on islands. While ROCK inhibitor, Y27632, did not significantly affect the behavior of Par-islands (data not shown), an actin inhibitor, Latrunculin B, changed the islands into a spherical shape, which frequently formed membrane protrusions (Fig. 5**g**, Supplementary movie 5), suggesting that the actin-membrane skeleton is necessary to balance the surface tension of Par-islands (see Discussion).

## Discussion

In this study, we reconstructed Par-complex-dependent cortical cell polarity induced by Par3 overexpression in non-polar S2 cells, using the *Gal4-UAS* system and the *Metallothionein* promoter for Par3 expression. Because this polarity requires endogenous Par6, aPKC, and Lgl, the reconstruction system reproduced the fundamental properties of Par-dependent polarization *in vivo*, at least in part. While there is no firm information regarding the Par3 protein level in polarized cells *in vivo*, the ratio of overexpressed Par3 protein level to endogenous Par3, in S2 cells, was estimated to be approximately 300-fold and 20-fold for the *Gal4-UAS* system and *Metallothionein* promoter, respectively (Supplementary Fig. 5).

### Temporal patterns of Par-complex aggregation

In our reconstruction system, cortical asymmetry began with the formation and growth of cortical dot-like structures, which were also reportedly associated with anterior localization of the Par-complex in *C. elegans* zygotes(Wang *et al*, 2017; Dickinson *et al*, 2017; Munro *et al*, 2004). Par-dots in the S2 cell system included all 3 Par complex components. Thus, these dots appear to be the common initial process of Par-complex cortical aggregation. The subsequent process of asymmetric localization proceeds in the form of Par-islands with amorphous and dynamic behavior. To our knowledge, this structure has not been reported in cortical Par-complex assembly in *C. elegans* or *Drosophila*. However, island-like structures were observed during asymmetric Par complex distribution in *Drosophila* neuroblasts, suggesting that Par-islands were not specific to this artificial system that used apolar S2 cells.

In *C. elegans*, asymmetric segregation of the Par-complex is driven by cortical flow and asymmetric contraction of the actomyosin meshwork(Munro *et al*, 2004; Motegi & Sugimoto, 2006). However, there was no indication that cortical flow was involved in the asymmetric clustering of the Par-complex in S2 cells. Furthermore, no directional movement towards the pole of polarization was observed. Interestingly, initial dot formation appeared to be biased towards the region opposite the cleavage point, where the centrosome also appeared to be located, which was consistent with a recent study on *Drosophila*(Loyer & Januschke, 2018; Januschke & Gonzalez, 2010; Jiang *et al*, 2015). Thus, the cleavage point and/or the centrosome may be a general positional cue for the initiation of Par-complex-dependent cell polarity. In this context, polarization process of the S2 cell system is likely to be cell-autonomous and dependent on the induction of polarity proteins, wherein the orientation of polarity appeared to be dependent on internal cue(s).

### The morphology and dynamics of Par-islands

Par-complex assembly at the cortex of S2 cells appears to stabilize the cell membrane because membrane filopodia extensively formed in areas where Par-islands were absent (Supplementary movie 2). Also, cell membrane curvature was higher where Par-islands were attached, compared with that of the surrounding areas (Fig. 5**g**). Membrane curvature may be determined by the balance between elasticity of the cortical cytoskeleton, the affinity of the Par-complex towards the cell membrane, and possibly the surface tension of the Par-island. Membrane affinity of the Par-complex is mediated by Par3 MAR, which interacts with phosphoinositides (PIPs) (Krahn *et al*, 2010)and/or by Par6-cdc42 interaction(Joberty *et al*, 2000). The convex shape of the Par-island and its higher membrane curvature reflects its relatively high surface tension. This is supported by the fact that disruption of the actin cytoskeleton by Latrunculin B treatment leads to a curled or spherical Par-island, inducing dynamic cell membrane protrusions. This phenomenon may be explained as follows; disruption of the cortical cytoskeleton leads to the loss of its elasticity, which had balanced the surface tension of the Par-island. The resulting imbalance in surface tension may cause the Par-island to shrink into a bowl or sphere shape, thereby bending the cell membrane outward and conferring protrusive activity to the cell membrane. In contrast, when membrane affinity is quite low, as in the case of Par3ΔMAR, Par-island shape is not affected by either cortical cytoskeleton elasticity or membrane affinity, and its shape would be determined only by the surface tension of Par3-islands. Under these conditions, we found that the Par-complex forms small cytoplasmic droplets, which subsequently coalesce into a spherical, densely packed structure, suggestinng that phase separation takes place between the Par-complex aggregates and the cytoplasm(Hyman *et al*, 2014).

### Molecular network of the Par-complex in the island state

In this study, we revealed that a Par-island is a meshwork of various polygonal shapes, which appear to be unit-like segments with an average length of approximately 0.4 μm. Isolated fragments such as single fragments and structures made up of a few connected fragments were observed during the development of Par-islands via live-imaging. These observations suggested that these isolated fragments assembled into a meshwork to form islands. These islands change shape rapidly during their movement along the cortex, and sometimes fuse to release pieces of different sizes, raising the possibility that Par-islands and small free fragments are mutually exchangeable. The factors that determine the size of these unit segments need further investigation.

Par3 is known to polymerize *in vitro* via the CR1 domain at its N-terminus to form a helical polymer of 8-fold symmetry(Zhang *et al*, 2013; Feng *et al*, 2007). Whether Par3 polymers are involved in the cortical cluster of the Par complex remain unclear. Our super-resolution microscopic observations and the ability of Par3 to form filaments lead to the simple hypothesis that the unit segment of a Par-island is formed by Par3 polymers as the core structure. Cell phenotypes expressing Par3 lacking its oligomerization domain, Par3ΔCR1, is compatible with this hypothesis. While there are many possibilities via which Par3 filaments may form a unit segment, a single Par3 polymer may form a single segment. Another possibility is for Par3 polymers to be aligned along the long axis of the segment. Since Par6 and aPKC bind the PDZ and CR3 domains of Par3, respectively, Par6 and aPKC may act as cross linkers between segments(Feng *et al*, 2007). Given the phenotype of ParS980A overexpression, the association of Par3 and aPKC by aPKC phosphorylation may confer flexibility and dynamism to the structure and/or assembly of the segmental elements. These hypotheses need to be tested in future studies.

### The two states of the Par-island distribution at steady state

An interesting property of Par-islands is that they are not unified into one large island under the cell membrane, even when polarized. Overexpression of Par3ΔMAR or Par3S980A is an exception. In the latter case, rapid and stable formation of the cortical Par-complex does not seem to permit separate island formation, and a large, transient dome is formed instead. In the former case, the Par-complex aggregates to form one large sphere. This cytoplasmic phenomenon is likely to be due to a phase separation between the Par-complex and the cytoplasm. Considering this property of the Par-complex, the unique feature of Par-islands associated with the cell membrane may reflect phase separation in 2 dimensions.

Steady state Par-island distribution in a cell may be classified into two different states, polarized and non-polarized. While we failed to identify a single parameter correlating these 2 states(Supplementary Fig. 2**b-d**), our analysis shows that the 2 states of island distribution are nearly fixed during the formation of islands (Supplementary Fig. 2**e**). Because the position of island formation appears to be stochastic, variation in the position of Par-island formation across the cell may explain the 2 localization patterns of Par-islands. Since Lgl distribution is largely complementary to dots and islands, this molecule may contribute to stabilize the 2 states of island distribution at the cellular scale(Betschinger *et al*, 2003; Guo & Kemphues, 1995). Thus, these 2 different states of Par-island distribution may be the outcome of 2 stable solutions of the reaction diffusion system(Chau *et al*, 2012; Goehring *et al*, 2011), where a negative regulator Lgl is involved(Betschinger *et al*, 2003). The initial condition, which is possibly determined by a stochastic distribution of islands, may select one of the two stable patterns in a cell. We propose that such cell-scale patterning is coupled with local phase separation of Par-islands as previously described for the membrane lipid domain(John & Bär, 2005).

In summary, we have developed a potential Par complex-polarization system upon induction of Par3 in non-polar S2 cells, which provides a useful model for cell-autonomous cell polarization, allowing us to easily manipulate gene expression and image at the super-resolution level. One intriguing challenge will be the coupling of mitosis with cell polarization in this system to induce asymmetric division.

## Materials and Methods

### Cell culture

S2 cell culture and transfection were performed as previously described (Ogawa et al., 2009). Expression vectors were transfected at two days prior to microscopic or Western blot analysis. For induction of the *Metallothionein* promoter, 100 mM CuSO_4_ solution was added to a medium at a final concentration of 1 mM.

### Live cell imaging

Cells were mounted on a 35 mm glass-bottom dish coated with 15 μg/ml poly-L-ornithine and incubated at 25°C for 30 min, followed by microscopic analysis. Images were taken at a 1 μm z-interval with a spinning disk confocal microscopy CSU-W1 (Yokogawa, Tokyo, Japan) equipped with a sCMOS camera Neo (Andor, Belfast, Northern Ireland) and MetaMorph software (Molecular Devices, San Jose, CA, USA).

### Immunostaining

For immunostaining of S2 cells, transfected cells were mounted on a poly-L-ornithine-coated cover slip and fixed with 4% paraformaldehyde in PBS for 15 min at room temperature. Cells were washed with PBS, followed by treatment with 0.1% Triton-X100 in PBS for 15 min. After washed with PBS, cells were treated with a blocking buffer containing 2% BSA in PBS for 30 min and incubated with primary antibodies in the blocking buffer for 30 min, followed by incubation with secondary antibodies for 30 min. Immunostained cells were embedded in mounting medium PermaFluor (Thermo Fisher Scientific) and analyzed with a confocal microscopes LSM510 (Zeiss, Oberkochen, Germany). For super-resolution microscopy, samples were embedded in ProLong Glass Andifade Mountant (Thermo Fisher Scientific).

For immunostaining of *Drosophila* neuroblasts, brains isolated from third instar larvae were fixed with 4% paraformaldehyde in PBS for 20 min at room temperature. Samples were treated with the blocking buffer for 1 h, followed by incubation with primary antibodies and secondary antibodies for 1 h each. Samples were then embedded in ProLong Gold Antifade Mountant (Thermo Fisher Scientific) and analyzed with a confocal microscope LSM880 (Zeiss).

Primary antibodies used were anti-aPKC (rabbit polyclonal, used at 1:1000, Santa Cruz), anti-Par-3 (rabbit polyclonal, used at 1:1000 or mouse monoclonal, used at 1:100)(Ohshiro *et al*, 2000), anti-Par-6 (rabbit polyclonal, used at 1:1000)(Izumi *et al*, 2004), anti-Myc (chicken polyclonal, used at 1:1000, Bethyl), anti-Flag (mouse monoclonal, used for 1:1000), anti-Miranda (Mouse monoclonal, used at 1:100(Ohshiro *et al*, 2000), anti-Lgl (rabbit polyclonal, used for 1:1000) (Ohshiro *et al*, 2000), anti-GFP (chicken polyclonal or mouse monoclonal, used for 1:1000). Secondary antibodies used were anti-Rabbit Alexa Fluor488 (Donkey polyclonal, Jackson Immuno Research), anti-Mouse Alexa Fluor488 (Donkey polyclonal, Jackson Immuno Research), anti-chicken Alexa Fluor488 (Donkey polyclonal, Jackson Immuno Research), anti-Rabbit Cyanin3 (Donkey polyclonal, Jackson Immuno Research), anti-Mouse Cyanin3 (Donkey polyclonal, Jackson Immuno Research), anti-chicken Cyanin3 (Donkey polyclonal,, Jackson Immuno Research), anti-Rabbit Alexa Fluor647 (Donkey polyclonal, Jackson Immuno Research). All of them are used at 1:4000.

### Super-resolution microscopy

For the super-resolution radial fluctuations (SRRF) methodc, confocal imaging was performed using LSM880 (Zeiss) with an objective lens Plan-Apochromat 63x/1.4 Oil DIC M27 (Zeiss). A series of 200 frames was obtained for each cell with a pixel size of 53 nm and 160 ms exposure time. Drift-correction and reconstruction of SRRF images were performed with an ImageJ plug-in NanoJ-SRRF(Gustafsson *et al*, 2016).

Using SRRF-processed images, Par3 contour lengths along the meshwork were manually traced with Fiji. Each image was overlayed by an edge-enhanced image generated with the Sobel filter, to highlight Par3 contour shapes. Lengths between their terminal ends and/or branching points were measured. A histogram and a density plot were generated from all contour lengths, and the shape of the density plot was fitted with a linear combination of 7 Gaussian curves by a fitting function implemented in R with the non-linear least square method. Power spectral density of the second derivative of the density plot was calculated using fast Fourier transform method with R.

Stimulated emission depletion (STED) imaging was performed using TCS SP8 STED 3X microscope (Leica, Wetzlar, Germany) with an objective lens HC PL APO 93X/1.30 GLYC (Leica). Deconvolution was performed with a deconvolution software package Huygens Professional (version 17.10, Scientific Volume Imaging, Hilversum, Netherlands).

Deconvoluted STED images were used for the analyses of Par3 segment lengths and widths. The segment length was defined as a shortest length between terminal ends, corners and/or branching points of Par3 contours, and manually traced with Fiji. The segment width was given by the full width at half maximum (FWHM) of a Gaussian-fitted signal distribution orthogonal to each Par3 segment.

### Quantification of asymmetry and statistics

The equatorial z-plane of each cell was analyzed for the estimation of asymmetric index (ASI) (see also Supplementary Fig. 1**b**). The cell perimeter was traced by a 0.5 μm-width line and the signal intensity along the line was measured with Fiji. The signal intensities were summed up along the half (L) of the total perimeter length (2L). The difference between this value and that of the other half was calculated and normalized by the total signal intensity along the perimeter. This measurement was done starting from every pixel along the perimeter (1 pixel = 0.108 μm), The maximum value of them was defined as ASI. ASI larger than 0.35 was defined as polarized cell, and the statistical significance of polarized cell population was analyzed by Fisher’s exact test with post-hoc Bonferroni correction for multiple comparisons (Fig. 3a, and Fig. 5e, -f). Statistical analyses were performed with R software.

### Western blot analysis

Whole cell extracts of the untransfected S2 cells and the transfected S2 cells were subjected to SDS-polyacrylamide gel electrophoresis. Primary antibodies used were anti-Par3 antibody (rabbit polyclonal, used at 1:1000), anti-alpha-tubulin (rat monoclonal, Santa Cruz). Secondary antibodies used were horseradish peroxidase (HRP)-conjugated anti-mouse antibody (sheep polyclonal, used at 1:3000, GE Healthcare), HRP-conjugated anti-rabbit antibody (sheep polyclonal, used at 1:3000, GE Healthcare) and HRP-conjugated anti-chicken antibody (donkey polyclonal, used at 1:250, SA1-300, Thermo Fisher Scientific). Protein level was analyzed by chemiluminescence with Chemi-Lumi One L (Nacalai tesque, Kyoto, Japan) and quantified with an image analyzer LAS-3000 system (Fujifilm, Tokyo, Japan). To compare the expression level of the overexpressed fluorescent protein per cell between two different transfectants or with the endogenous Par3 proteins, transfection efficiency for each sample was calculated by counting fluorescence-positive cells and negative cells. The ratio of the expression level per cell was calculated by dividing the measured staining intensity on the Western blot by the transfection efficiency.

### Knock-down experiment

Long double-stranded RNAs (dsRNAs) were used for RNA interference (RNAi) in S2 cells as previously described(Bettencourt-Dias & Goshima, 2009). dsRNA for knocking-down *Par-6* or *aPKC* was synthesized with MEGAscript T7 Transcription Kit (Ambion, Thermo Fisher Scientific) according to the manufacurer’s instructions, by using *pBS-T7/Par-6/T7* or *pBS-T7/aPKC/T7* plasmid, which contains a full-length ORF of Par-6 or aPKC flanked by two T7 promoters, as a template, respectively. dsRNA for knocking-down Lgl was by using *pUAS-Flag-Lgl*, and primers in below. Forward:

5’-TAATACGACTCACTATAGGGATGGCAATAGGGACGCAAACAGGGGCTTTAAAAGTT-3’, Reverse:

5’-AATACGACTCACTATAGGGTTAAAATTGGCTTTCTTCAGGCGCTGTTTTTGGCGTTCCAA-3’. dsRNAs were added to the culture media at a final concentration of 4.5 μg/ml at 2–3 h following transfection of expression plasmids.

### Plasmid construction

To construct expression vectors under control of an *actin (act5c)* promoter, *Drosophila Par6, Par3 or aPKC* ORFs, or *Par6, aPKC* and myristoylation tag from *Fyn* (5’-ATGGGCTGTGTGCAATGTAAGGATAAAGAAGCAACAAAACTGACG-3’) conjugated with *GFP* at the C-terminus (*Par-6-GFP* and *aPKC-GFP*) was inserted into *pAc5.1/V5-His B* plasmid (Invitrogen, Thermo Fisher Scientific). To construct an expression vector for Par-3 under control of the *Gal4-UAS* system, *Drosophila gal4* ORF was inserted into *pDAMCS* plasmid. To constructed *pDAMCS* plasmid, Bgl **II**-Xho **I** fragment of *pAct5C0* plasmid(Thummel *et al*, 1988) containing actin 5C promoter and poly(A) addition signals (and a small region of hsp70 promoter) was cloned between BamH **I** and Sal **I** site of *pUC19* plasmid. Then, a synthetic double-strand oligonucleotide containing multiple cloning sites was cloned into the BamH **I** site to produce pDAMCS expression plasmid. Par-3 conjugated with *Myc* or *Flag* epitope and *mKate2* at the N-terminus and C-terminus, respectively, and *Lgl*^*3A*^ with *Flag* epitope was inserted into *pUAST* plasmid(Brand & Perrimon, 1993). To construct expression vectors for Par-3 under the control of the induction system, Par-3 that had been conjugated with *Myc* epitope and mKate2 at the N-terminus and C-terminus, respectively, was inserted into *pMT* plasmid (Invitrogen, Thermo Fisher Scientific). pUbq-Spd2-GFP is kindly provided by Jordan Raff (University of Oxford, UK). Plasmids used for transfection were purified with Wizard Plus SV Minipreps DNA Purification System (Promega, Madison, WI, USA) or NucleoBond Xtra Midi (Macherey-Nagel, Düren, Germany).

## Acknowledgements

We thank Y. Tsunekawa for technical advice, J. Raff for plasmids, T. Nishimura for fly strains, S. Hayashi, T. Nishimura, and K. Kawaguchi for critical discussion. Some of the imaging experiments were performed at the RIKEN Kobe Light Microscopy Facility. I.F. is a recipient of RIKEN Special Postdoctoral Researcher Program.

## Contributions

K.K., S.Y., I.F. and F.M. designed the project. K.K. engaged in and performed all the experiments. K.K, I.F., A.S. and Y.O performed super resolution microscopic observations. K.K., S.Y., A.S., I.F. and F.M. analyzed data. K.K., I.F., T.S. and F.M. prepared the manuscript with input from all authors.

## Data availability

All raw data are available upon the request.

## Supplementary Figure legends

**Supplementary Fig. 1. Par-polarity induction by the Gal4>UAS-Par3**

**a.** Comparison of the expression level of Par3-GFP driven by the *actin* promoter with that driven by the *actin*-promoter-*Gal4* × *UAS* system. Western blotting was performed for S2 cells transfected with *pAct-Par3-GFP* (100 μg and 300 μg /10^6^ cells) and with *pAct-Gal4* and *pUAS-Par3-GFP*, and the blot was stained with the anti-Par3 antibody.

**b.** Definition of the asymmetric index (ASI). ASI is defined by the maximum difference in the cumulative intensity of fluorescence (such as Par3-mKate2 and Par6-GFP) between a half cell perimeter and the other half at the equatorial plane of the cell, normalized via dividing by the cumulative intensity of the entire cell perimeter.

**c.** ASI distribution was compared between cells expressing memGFP and those expressing Par3-mKate2 at the cell cortex, both of which were driven by the Act-Gal4xUAS system. ASI value of memGFP is distributed broadly and ranges from 0 to 0.35 (mean value = 0.14±0.08). Since memGFP, in principle, has no ability to polarize, such a wide distribution originated from the fluctuation of random distribution along the equatorial cell perimeter and also from the existence of local membrane flairs. Distribution of ASI for cells showing cortical Par3 distribution (mean = 0.34±0.18) may be categorized into 2 groups (Fig. 2C); a group of cells show low ASIs overlapping with those of cells expressing memGFP in the range from 0 to 0.35 (approximately 62% of the transfected cells that localize mKate2 cortically), indicating that cells belonging to this group are essentially non-polarized. The ASI values of the other group (approximately 38%), are broadly distribute, but display ASI values larger than the ASI distribution of mem-GFP cells (> 0.35, mean value = 0.52±0.14)

**Supplementary Fig. 2. Temporal pattern and steady state of the clustering of Par-islands**

**a.** Temporal pattern of the fluorescence intensity of Par6-GFP expressed in S2 cells transfected with *pAct-Par6-GFP*. The fluorescence intensity of Par6-GFP was measured for 6 cells every 1 h from two days following transfection. Expression levels did not drastically change from 6 h onward following transfection, indicating that Par6-GFP is an appropriate marker for the Par complex distribution in cells, when Par3-mKate2 was induced.

**b-d.** Temporal pattern of the fluorescence intensity of Par3-mKate2 (**b**), its rate of change (**c**) and the ratio of Par6-GFP/Par3-mKate2 (**d**), and the ASI value (**e**) of S2 cells that were transfected with *pMT-Par3-mKate2* and *pAct-Par6-GFP*, followed by induction by CuSO4 addition two days after plasmid transfection. Time 0 is the timing of CuSO4 addition (two days following plasmid transfection). Measurements were taken every 30 min. Fluorescence intensity reached the steady level around 8 h after induction (**b,e**). In (**b-e**), the blue line indicates the averaged values of 10 cells that showed a non-polarized distribution of Par6-GFP at 8 h after induction (with the ASI value around 0.2). The red line indicates the average values of 13 cells that showed polarized distribution of Par6-GFP following 8 h induction (with the ASI value around 0.4). Bars indicate s.d. **b-d**. The temporal pattern of the fluorescence intensity/cell of Par3-mKate2 and Par6-GFP/Par3-mKate2 ratio are not significantly different between the polarized cell group (red line) and non-polarized cell group (blue line). **e**. ASI value began to increase immediately after the rise of Par3-mKate2 levels (approximately 2 h after induction) in the polarized cells (blue line), and maintained a high level afterwards. On the other hand, the non-polarized cells (red line) initially showed a slight increase in the ASI value, but decreased from 5 h after induction onwards. The timing of the increase in ASI roughly corresponded to that of Par-dot emergence (2-4 h after induction; see Fig. 2**a-c**), and the timing of a decrease in ASI value in non-polarized cells roughly corresponds to the late period of Par-island formation (4-6 h after induction), although there are cell to cell variations in these timings. *t* test, *P* = 5.6×10^−5^, 0.04, 5.5×10^−4^, 1.1×10^−6^, 5.5×10^−4^, and 4.3×10^−7^ for every 30 min time point from 6 h after induction.

**f.** Comparison of Par3-mKate2 expression level induced by the *Metallothionein* promoter, *pMT-Par3-mKate2*, and that promoted by the *pAct-Gal4xUAS* system. The mKate2 fluorescence intensity of the individual S2 cells was measured two days after transfection of *pAct-Gal4 and UAS-Par3-mKate2*, or at 8 h post CuSO_4_ induction of *pMT-Par3-mKate2*, two days after transfection of the plasmid. The expression level of Par3-mKate2 per cell was approximately 18-fold higher when it was driven by the *UAS-Gal4* system (6.1×10^4^ ±7.1×10^4^) than that of the steady state level induced by the *Metallothionein* promoter (3.4×10^3^ ±9.8x 10^2^).

**g.** Stability of polarized and non-polarized cells. ASI values 11 h post-induction were measured for cells polarized 8 h post-induction (ASI>0.4, n=11) and for non-polarized cells (ASI < 0.3, n=14). ASI values were measured using induced -Par6-GFP.

**h-j**. Relationship between the Par-complex crescent and the position of the centrosome. The centrosome was visualized via Spd-GFP, which was expressed by the transfection of *pUbq-Spd-GFP* and *pDA-Gal4* together with *pUAS-Par3*. Spd2-GFP and aPKC were immunostained (**h**). The angle between the radial lines from the cell center to 2 edges of the aPKC crescent (θ1), and the angle between an edge of the crescent and the centrosome (θ2) in the clock-wise direction were measured at the equatorial plane (**i**). In 20 out of 27 cells (74%), the centrosome was located within the fan shape made by the aPKC crescent and the cell center (**j**).

**Supplementary Fig. 3. SRRF analysis of Par-island meshwork**

**a.** Left. the average of 200 confocal images of Par3-mKate2 distribution in a cell that expresses Par3-mKate2 and Par6-GFP. Those 200 images were used for SRRF analysis (Fig. 4**a)**. The middle Panel shows the SRRF-processed image of Fig. 4**a**, which was processed with edge detection (see Methods). By this process, the continuous contours become clearly visible. Edges in the image were visualized in green. The right panel shows tracing of continuous contours of the SRRF-processed image (Fig. 4**a)** after edge detection (light blue lines). The distribution of the lengths of the continuous lines is shown as a histogram in Fig. 4**c**. Scale bar, 5 μm.

**b.** The list of means and s.d., of 7 Gaussian curve (Fig. 4**d**), combinations that best fit the density plot of the continuous contour lengths distribution shown in Fig. 4**d** (see Methods).

**Supplementary Fig. 4 STED analysis of Par-island meshwork**

**a.** STED image of a cell that expresses Par3-GFP. The distribution of GFP was detected via indirect immunofluorescence staining. The image on the left in Fig. 4**e** shows the deconvolved image (see Methods). Scale bar, 10 μm for (**a-d**).

**b.** Individual segments composing Par-islands were traced in the image (**a**) after deconvolution (Fig. 4**e**). Tracing of the segments is indicated as light blue straight lines.

**c.** STED image of a cell expressing Par3-GFP. The deconvolution of the original image is shown on the right in Fig. 4**e**.

**d.** Tracing of individual segments of Par-islands in the right-side image of Fig. 4**e**. The lengths of segmental rods detected in these 2 images (**b,d**) are plotted in Fig. 4**h**.

**e.** Distribution of half widths of segments composing Par-islands that were visualized by GFP staining in the two cells shown in Fig. 4a. The mean is 0.22±0.03 μm (n=19).

**f.** An example of Gaussian fitting of the fluorescence intensity distribution along the segment width visualized via immunofluorescence-staining for GFP (half width=0.24 μm).

**g.** Rod and string structures of the Par3-mKate2 in 3D time-lapse images of a cell expressing Par3-mKate2 during the period of Par-dot formation and development. Four time points were selected from Supplementary movie 2. Scale bar, 2 μm. Insets display the magnification of a part of the image. Arrowheads indicate Par-dots, arrows, rods, and strings. Scale bar, 0.5 μm.

**Supplementary Fig. 5. Comparison of Par3 expression level**

Comparison of expression levels between Par3 induced by the *MT* promoter and endogenous Par3 determined via Western blotting. S2 cells were transfected with *pMT-myc-Par3-mKate2*, and induced for 8 h using CuSO_4_ from two days post-transfection for Western blotting. Left lane: 8 h-induction of myc-Par3-mKate2 by the *MT* promoter, right lane: not transfected with *pMT-myc-Par3-mKate2*.

### List of movies

Movie 1: 3D time-lapse movie of a polarized S2 cell monitored by Par6-GFP two days after transfection of pAct-Gal4 > UAS-myc-Par3-mKate2, pAct-Par6-GFP, and pAct-aPKC. Par-islands are clustered with dynamic movements.

Movie 2: 3D time-lapse movie of a S2 cell monitored by Par6-GFP following induction of myc-Par3-mKate2 from *Metallothionein* promoter. Induction started at time 0 by the addition of CuSO4 two days after transfection of pMT-myc-Par3-mKate2, pAct-Par6-GFP, and pAct-aPKC.

Movie 3: 3D time-lapse movie of a polarized S2 cell monitored by Par6-GFP at 8 h induction of myc-Par3-mKate2 with the co-expression of pAct-Par6-GFP and pAct-aPKC.

Movie 4: 3D movie of a polarized S2 cell stained for myc-Par3-mKate2 and Lgl at 8 h induction of myc-Par3-mKate2 with the co-expression of pAct-Par6-GFP and pAct-aPKC.

Movie 5: 3D movie of a nonpolarized S2 cell stained for myc-Par3-mKate2 and Lgl at 8 h induction of myc-Par3-mKate2 with the co-expression of pAct-Par6-GFP and pAct-aPKC. A part of the adjacent cell is included in the movie.

Movie 6: 3D time-lapse movie of a S2 cell monitored by Par6-GFP, 3 min after the addition of Latrunculin B, at 8 h induction of myc-Par3-mKate2, following two days transfection of pMT-myc-Par3-mKate2, pAct-Par6-GFP, and pAct-aPKC.

## References

Aceto D, Beers M & Kemphues KJ (2006) Interaction of PAR-6 with CDC-42 is required for maintenance but not establishment of PAR asymmetry in *C. elegans*. Developmental Biology 299: 386–397

Baas AF, Kuipers J, van der Wel NN, Batlle E, Koerten HK, Peters PJ & Clevers HC (2004) Complete polarization of single intestinal epithelial cells upon activation of LKB1 by STRAD. Cell 116: 457–466

Benton DSJR (2003) A Conserved Oligomerization Domain in *Drosophila* Bazooka/PAR-3 Is Important for Apical Localization and Epithelial Polarity. 13: 1317–1323

Betschinger J, Mechtler K & Knoblich JA (2003) The Par complex directs asymmetric cell division by phosphorylating the cytoskeletal protein Lgl. Nature 422: 326–330

Bettencourt-Dias M & Goshima G (2009) RNAi in *Drosophila* S2 cells as a tool for studying cell cycle progression. Methods Mol. Biol. 545: 39–62

Brand AH & Perrimon N (1993) Targeted gene expression as a means of altering cell fates and generating dominant phenotypes. Development 118: 401–415

Chau AH, Walter JM, Gerardin J, Tang C & Lim WA (2012) Designing synthetic regulatory networks capable of self-organizing cell polarization. Cell 151: 320–332

Derivery E, Seum C, Daeden A, Loubéry S, Holtzer L, Jülicher F & Gonzalez-Gaitan M (2015) Polarized endosome dynamics by spindle asymmetry during asymmetric cell division. Nature 528: 280–285

Dickinson DJ, Schwager F, Pintard L, Gotta M & Goldstein B (2017) A Single-Cell Biochemistry Approach Reveals PAR Complex Dynamics during Cell Polarization. Developmental Cell 42: 416–434.e11

Doerflinger H, Benton R, Torres IL, Zwart MF & St Johnston D (2006) *Drosophila* anterior-posterior polarity requires actin-dependent PAR-1 recruitment to the oocyte posterior. Current Biology 16: 1090–1095

Feng W, Wu H, Chan LN & Zhang M (2007) The Par‐3 NTD adopts a PB1‐like structure required for Par‐3 oligomerization and membrane localization. The EMBO Journal 26: 2786–2796 Available at: http://emboj.embopress.org/content/26/11/2786.abstract

Goehring NW, Trong PK, Bois JS, Chowdhury D & Grill SW (2011) Polarization of PAR Proteins by Advective Triggering of a Pattern-Forming System. Science 403:

Guo S & Kemphues KJ (1995) pa3r-1, a Gene Required for Establishing Polarity in *C. elegans* Embryos, Encodes a Putative Ser / Thr Kinase That Is Asymmetrically Distributed. Cell 81: 611–620

Gustafsson N, Culley S, Ashdown G, Owen DM, Pereira PM & Henriques R (2016) Fast live-cell conventional fluorophore nanoscopy with ImageJ through super-resolution radial fluctuations. Nature Communications 7: 12471

Harris TJC (2017) Protein clustering for cell polarity: Par-3 as a paradigm. F1000Res 6: 1620

Hurov JB, Watkins JL & Piwnica-Worms H (2004) Atypical PKC phosphorylates PAR-1 kinases to regulate localization and activity. Current Biology 14: 736–741

Hyman AA, Weber CA & Jülicher F (2014) Liquid-Liquid Phase Separation in Biology. Annu. Rev. Cell Dev. Biol. 30: 39–58

Ikeshima-Kataoka H, Skeath JB, Nabeshima Y-I, Doe CQ & Matsuzaki F (1997) Miranda directs Prospero to a daughter cell during Drosophila asymmetric divisions. Nature 390: 625–629

Izumi Y, Hirose T, Tamai Y, Hirai S, Nagashima Y, Fujimoto T, Tabuse Y, Kemphues KJ & Ohno S (1998) An atypical PKC directly associates and colocalizes at the epithelial tight junction with ASIP, a mammalian homologue of *Caenorhabditis elegans* polarity protein PAR-3. J Cell Biol 143: 95–106

Izumi Y, Ohta N, Itoh-Furuya A, Fuse N & Matsuzaki F (2004) Differential functions of G protein and Baz-aPKC signaling pathways in *Drosophila* neuroblast asymmetric division. J Cell Biol 164: 729–738

Januschke J & Gonzalez C (2010) The interphase microtubule aster is a determinant of asymmetric division orientation in *Drosophila* neuroblasts. The Journal of cell biology 188: 693–706

Jiang T, McKinley RFA, McGill MA, Angers S & Harris TJC (2015) A Par-1-Par-3-Centrosome Cell Polarity Pathway and Its Tuning for Isotropic Cell Adhesion. Current Biology 25: 2701–2708

Joberty G, Petersen C, Gao L & Macara IG (2000) The cell-polarity protein Par6 links Par3 and atypical protein kinase C to Cdc42. Nat Cell Biol 2: 531–539

John K & Bär M (2005) Travelling lipid domains in a dynamic model for protein-induced pattern formation in biomembranes. Phys Biol 2: 123–132

Johnston CA, Hirono K, Prehoda KE & Doe CQ (2009) Identification of an Aurora-A/PinsLINKER/ Dlg Spindle Orientation Pathway using Induced Cell Polarity in S2 Cells. Cell 138: 1150–1163

Johnston DS (2018) Establishing and transducing cell polarity: common themes and variations. Current Opinion in Cell Biology 51: 33–41

Kemphues KJ, Priess JR, Morton DG & Cheng NS (1988) Identification of genes required for cytoplasmic localization in early C. elegans embryos. Cell 52: 311–320

Krahn MP, Klopfenstein DR, Fischer N & Wodarz A (2010) Membrane Targeting of Bazooka/PAR-3 Is Mediated by Direct Binding to Phosphoinositide Lipids. Current Biology 20: 636–642

Lang CF & Munro E (2017) The PAR proteins: from molecular circuits to dynamic self-stabilizing cell polarity. Development 144: 3405–3416

Loyer N & Januschke J (2018) The last-born daughter cell contributes to division orientation of *Drosophila* larval neuroblasts. Nature Communications 9: 1–12

Mizuno K, Suzuki A, Hirose T, Kitamura K, Kutsuzawa K, Futaki M, Amano Y & Ohno S (2003) Self-association of PAR-3-mediated by the conserved N-terminal domain contributes to the development of epithelial tight junctions. J. Biol. Chem. 278: 31240–31250

Morais-de-Sá E, Mirouse V & St Johnston D (2010) aPKC Phosphorylation of Bazooka Defines the Apical/Lateral Border in *Drosophila* Epithelial Cells. Cell 141: 509–523

Motegi F & Sugimoto A (2006) Sequential functioning of the ECT-2 RhoGEF, RHO-1 and CDC-42 establishes cell polarity in *Caenorhabditis elegans* embryos. Nat Cell Biol 8: 978–985

Munro E, Nance J & Priess JR (2004) Cortical flows powered by asymmetrical contraction transport PAR proteins to establish and maintain anterior-posterior polarity in the early *C. elegans* embryo. Developmental Cell 7: 413–424

Ohshiro T, Yagami T, Zhang C & Matsuzaki F (2000) Role of cortical tumour-suppressor proteins in asymmetric division of *Drosophila* neuroblast. Nature 408: 593–596

Plant PJ, Fawcett JP, Lin DCC, Holdorf AD, Binns K, Kulkarni S & Pawson T (2003) A polarity complex of mPar-6 and atypical PKC binds, phosphorylates and regulates mammalian Lgl. Nat Cell Biol 5: 301–308

Renschler FA, Bruekner SR, Salomon PL, Mukherjee A, Kullmann L, Schütz-Stoffregen MC, Henzler C, Pawson T, Krahn MP & Wiesner S (2018) Structural basis for the interaction between the cell polarity proteins Par3 and Par6. Sci. Signal. 11: eaam9899

Rodriguez J, Peglion F, Martin J, Hubatsch L, Reich J, Hirani N, Gubieda AG, Roffey J, Fernandes AR, St Johnston D, Ahringer J & Goehring NW (2017) aPKC Cycles between Functionally Distinct PAR Protein Assemblies to Drive Cell Polarity. Developmental Cell: 1–16

Rodriguez-Boulan E & Macara IG (2014) Organization and execution of the epithelial polarity programme. Nat Rev Mol Cell Biol 15: 225–242

Sailer A, Anneken A, Li Y, Lee S & Munro E (2015) Dynamic Opposition of Clustered Proteins Stabilizes Cortical Polarity in the C. elegans Zygote. Developmental Cell 35: 131–142

Schneider I (1972) Cell lines derived from late embryonic stages of *Drosophila* melanogaster. J Embryol Exp Morphol 27: 353–365

Soriano EV, Ivanova ME, Fletcher G, Riou P, Knowles PP, Barnouin K, Purkiss A, Kostelecky B, Saiu P, Linch M, Elbediwy A, Kjær S, O’Reilly N, Snijders AP, Parker PJ, Thompson BJ & McDonald NQ (2016) aPKC Inhibition by Par3 CR3 Flanking Regions Controls Substrate Access and Underpins Apical-Junctional Polarization. Developmental Cell 38: 384–398

Suzuki A & Ohno S (2006) The PAR-aPKC system: lessons in polarity. Journal of Cell Science 119: 979–987

Tabuse Y, Izumi Y, Piano F, Kemphues KJ, Miwa J & Ohno S (1998) Atypical protein kinase C cooperates with PAR-3 to establish embryonic polarity in *Caenorhabditis elegans*. Development (Cambridge, England) 125: 3607–3614

Thummel CS, Boulet AM & Lipshitz HD (1988) Vectors for Drosophila P-element-mediated transformation and tissue culture transfection. Gene 74: 445–456

Wang S-C, Yu T, Low F, Nishimura Y, Gole L, Yu W & Motegi F (2017) Cortical forces and CDC-42 control clustering of PAR proteins for *Caenorhabditis elegans* embryonic polarization. 19:

Yamanaka T, Horikoshi Y, Sugiyama Y, Ishiyama C, Suzuki A, Hirose T, Iwamatsu A, Shinohara A & Ohno S (2003) Mammalian Lgl Forms a Protein Complex with PAR-6 and aPKC Independently of PAR-3 to Regulate Epithelial Cell Polarity. Current Biology 13: 734–743

Yamashita YM, Yuan H, Cheng J & Hunt AJ (2010) Polarity in Stem Cell Division: Asymmetric Stem Cell Division in Tissue Homeostasis.: 1–14

Zhang Y, Wang W, Chen J, Zhang K, Gao F, Gao B, Zhang S, Dong M, Besenbacher F, Gong W, Zhang M, Sun F & Feng W (2013) Structural Insights into the Intrinsic Self-Assembly of Par-3 N-Terminal Domain. Structure 21: 997–1006

