## Supplemental Figures for "Reconstruction of Par polarity in apolar cells reveals a dynamic process of cortical polarization"

#### Supplementary Fig. 1

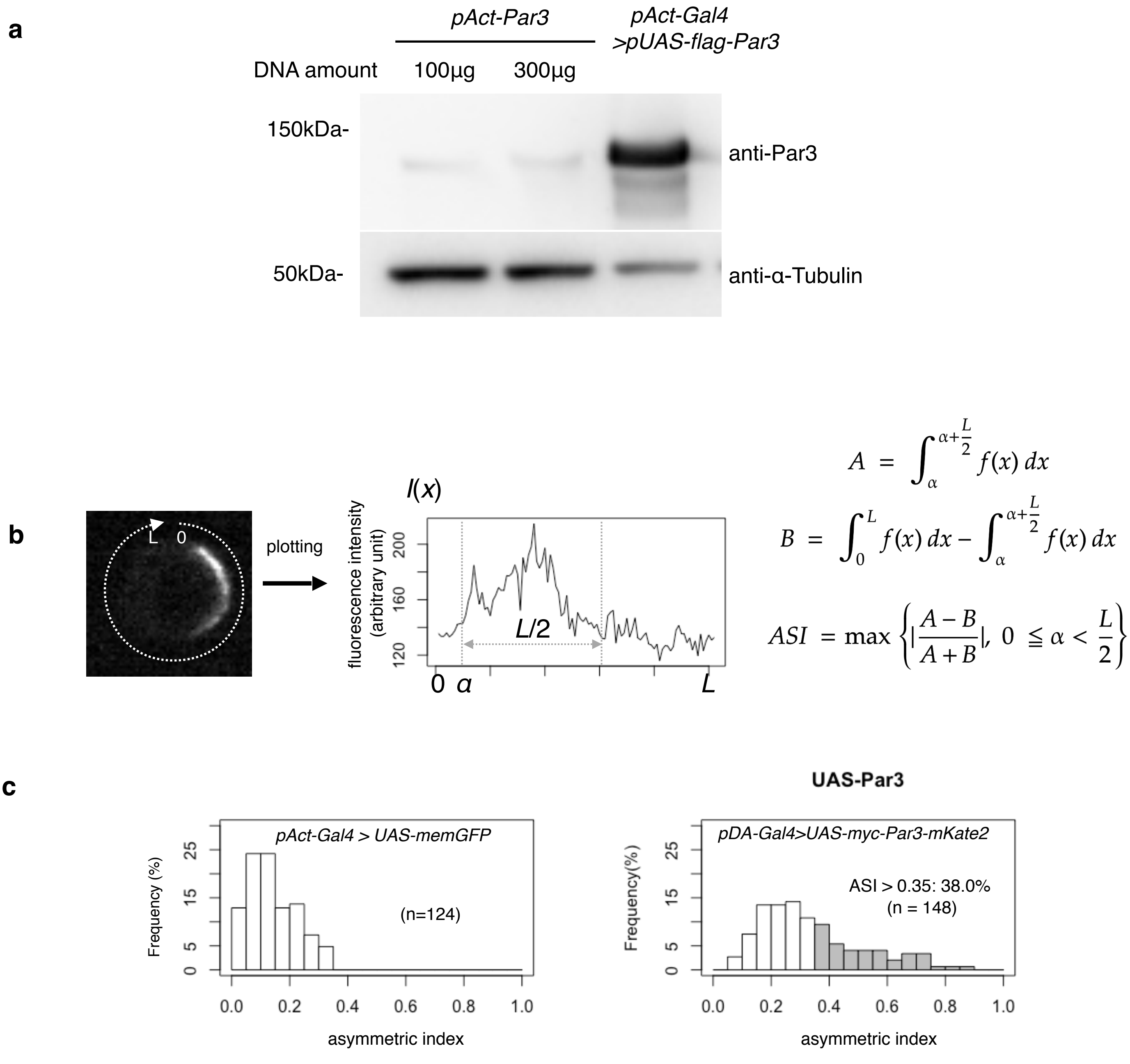

#### Supplementary Figure legends

**Supplementary Fig. 1. Par-polarity induction by the Gal4>UAS-Par3**

**a.** Comparison of the expression level of Par3-GFP driven by the *actin* promoter with that driven by the *actin*-promoter-*Gal4* x *UAS* system. Western blotting was performed for S2 cells transfected with *pAct-Par3-GFP* (100 µg and 300 µg /$10^6$ cells) and with *pAct-Gal4* and *pUAS-Par3-GFP*, and the blot was stained with the anti-Par3 antibody.

**b.** Definition of the asymmetric index (ASI). ASI is defined by the maximum difference in the cumulative intensity of fluorescence (such as Par3-mKate2 and Par6-GFP) between a half cell perimeter and the other half at the equatorial plane of the cell, normalized via dividing by the cumulative intensity of the entire cell perimeter.

**c.** ASI distribution was compared between cells expressing memGFP and those expressing Par3-mKate2 at the cell cortex, both of which were driven by the Act-Gal4xUAS system. ASI value of memGFP is distributed broadly and ranges from 0 to 0.35 (mean value = 0.14±0.08). Since memGFP, in principle, has no ability to polarize, such a wide distribution originated from the fluctuation of random distribution along the equatorial cell perimeter and also from the existence of local membrane flairs. Distribution of ASI for cells showing cortical Par3 distribution (mean = 0.34±0.18) may be categorized into 2 groups (Fig. 2C); a group of cells show low ASIs overlapping with those of cells expressing memGFP in the range from 0 to 0.35 (approximately 62% of the transfected cells that localize mKate2 cortically), indicating that cells belonging to this group are essentially non-polarized. The ASI values of the other group (approximately 38%), are broadly distribute, but display ASI values larger than the ASI distribution of mem-GFP cells (> 0.35, mean value = 0.52±0.14)

### Supplementary Fig. 2

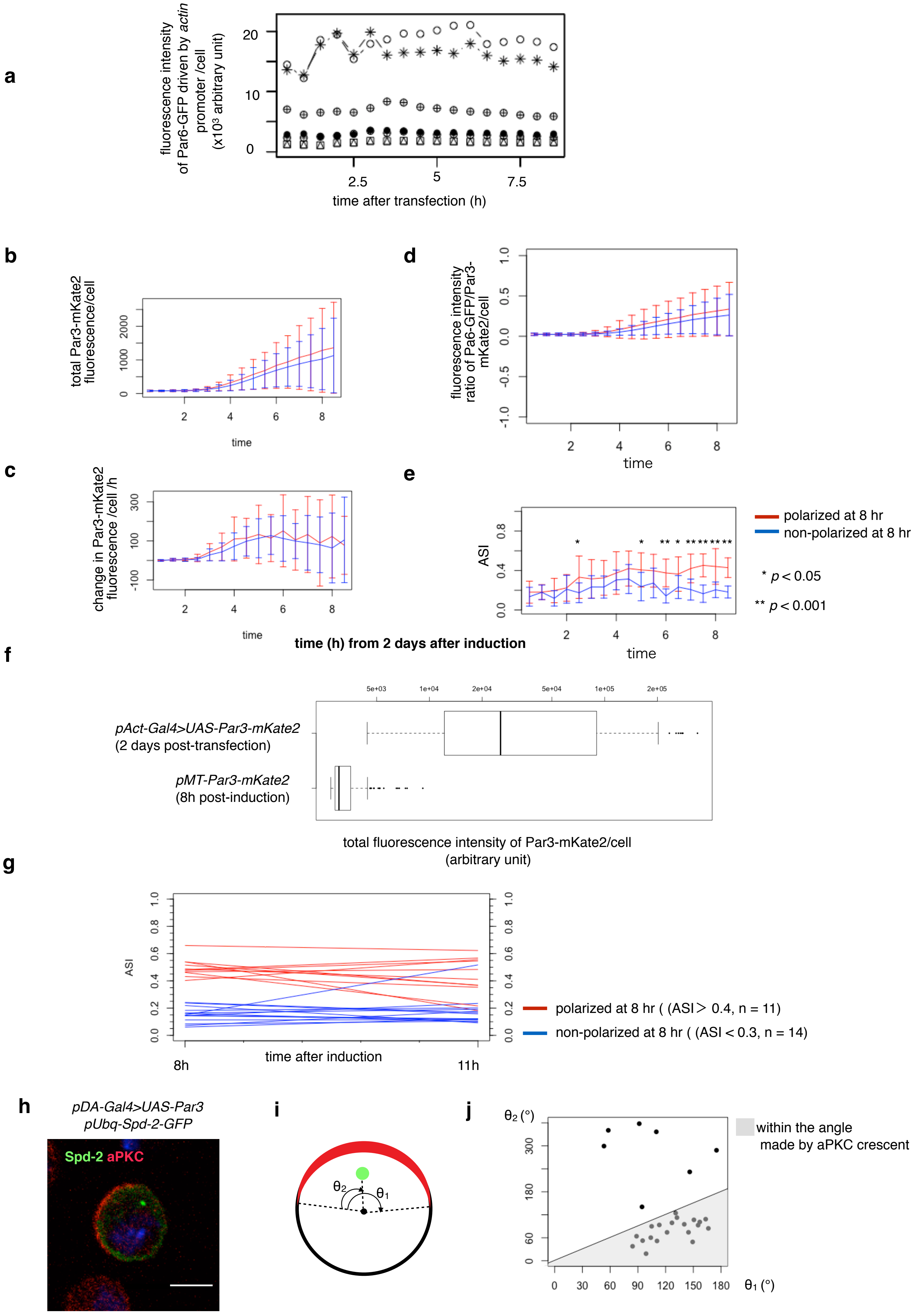

**Supplementary Fig. 2. Temporal pattern and steady state of the clustering of Par-islands**

**a.** Temporal pattern of the fluorescence intensity of Par6-GFP expressed in S2 cells transfected with *pAct-Par6-GFP*. The fluorescence intensity of Par6-GFP was measured for 6 cells every 1 h from two days following transfection. Expression levels did not drastically change from 6 h onward following transfection, indicating that Par6-GFP is an appropriate marker for the Par complex distribution in cells, when Par3-mKate2 was induced.

**b-d.** Temporal pattern of the fluorescence intensity of Par3-mKate2 (**b**), its rate of change (**c**) and the ratio of Par6-GFP/Par3-mKate2 (**d**), and the ASI value (**e**) of S2 cells that were transfected with *pMT-Par3-mKate2* and *pAct-Par6-GFP*, followed by induction by CuSO4 addition two days after plasmid transfection. Time 0 is the timing of CuSO4 addition (two days following plasmid transfection). Measurements were taken every 30 min. Fluorescence intensity reached the steady level around 8 h after induction (**b**,**e**). In (**b-e**), the blue line indicates the averaged values of 10 cells that showed a non-polarized distribution of Par6-GFP at 8 h after induction (with the ASI value around 0.2). The red line indicates the average values of 13 cells that showed polarized distribution of Par6-GFP following 8 h induction (with the ASI value around 0.4). Bars indicate s.d. **b-d**. The temporal pattern of the fluorescence intensity/cell of Par3-mKate2 and Par6-GFP/Par3-mKate2 ratio are not significantly different between the polarized cell group (red line) and non-polarized cell group (blue line). **e**. ASI value began to increase immediately after the rise of Par3-mKate2 levels (approximately 2 h after induction) in the polarized cells (blue line), and maintained a high level afterwards. On the other hand, the non-polarized cells (red line) initially showed a slight increase in the ASI value, but decreased from 5 h after induction onwards. The timing of the increase in ASI roughly corresponded to that of Par-dot emergence (2-4 h after induction; see Fig. 2**a-c**), and the timing of a decrease in ASI value in non-polarized cells roughly corresponds to the late period of Par-island formation (4-6 h after induction), although there are cell to cell variations in these timings. *t* test, $P = 5.6 \times 10^{-5}$, 0.04, $5.5 \times 10^{-4}$, $1.1 \times 10^{-6}$, $5.5 \times 10^{-4}$, and $4.3 \times 10^{-7}$ for every 30 min time point from 6 h after induction.

**f.** Comparison of Par3-mKate2 expression level induced by the *Metallothionein* promoter, *pMT-Par3-mKate2,* and that promoted by the *pAct-Gal4xUAS* system. The mKate2 fluorescence intensity of the individual S2 cells was measured two days after transfection of *pAct-Gal4 and UAS-Par3-mKate2*, or at 8 h post $CuSO_4$ induction of *pMT-Par3-mKate2,* two days after transfection of the plasmid. The expression level of Par3-mKate2 per cell was approximately 18-fold higher when it was driven by the *UAS-Gal4* system ($6.1 \times 10^4 \pm 7.1 \times 10^4$) than that of the steady state level induced by the *Metallothionein* promoter ($3.4 \times 10^3 \pm 9.8 \times 10^2$).

**g.** Stability of polarized and non-polarized cells. ASI values 11 h post-induction were measured for cells polarized 8 h post-induction (ASI>0.4, n=11) and for non-polarized cells (ASI < 0.3, n=14). ASI values were measured using induced -Par6-GFP.

**h-j**. Relationship between the Par-complex crescent and the position of the centrosome. The centrosome was visualized via Spd-GFP, which was expressed by the transfection of *pUbq-Spd-GFP* and *pDA-Gal4* together with *pUAS-Par3*. Spd2-GFP and aPKC were immunostained (**h**). The angle between the radial lines from the cell center to 2 edges of the aPKC crescent (θ1), and the angle between an edge of the crescent and the centrosome (θ2) in the clock-wise direction were measured at the equatorial plane (**i**). In 20 out of 27 cells (74%), the centrosome was located within the fan shape made by the aPKC crescent and the cell center (**j**).

**Supplementary Fig. 3**

**a**

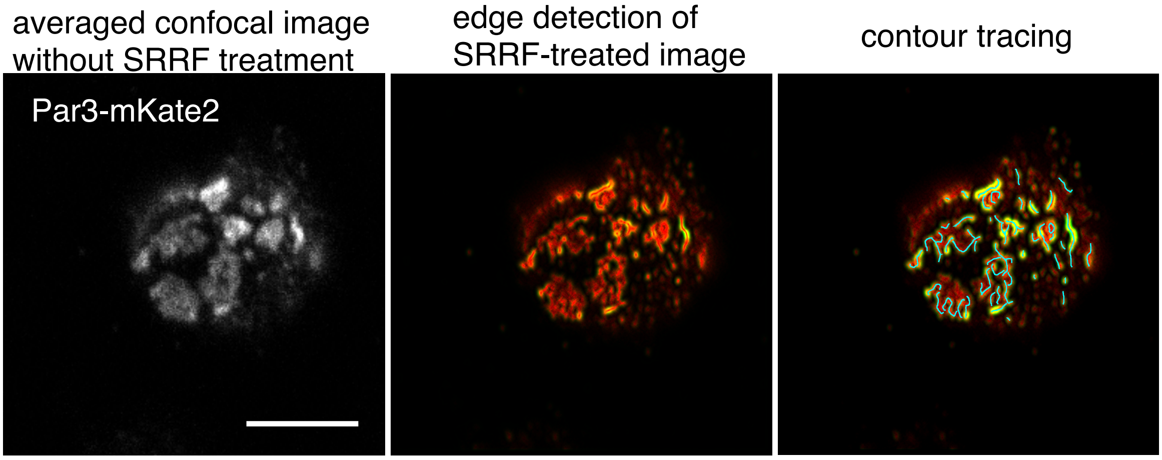


**b**

| peak # | 1 | 2 | 3 | 4 | 5 | 6 | 7 |
|---|---|---|---|---|---|---|---|
| mean | 0.27 | 0.64 | 1.10 | 1.48 | 1.92 | 2.26 | 2.63 |
| s.d. | 0.11 | 0.23 | 0.16 | 0.15 | 0.12 | 0.12 | 0.12 |
| weight | 0.24 | 0.46 | 0.13 | 0.09 | 0.04 | 0.02 | 0.02 |
| interval | 0.27 | 0.37 | 0.45 | 0.39 | 0.44 | 0.34 | 0.36 |

**Supplementary Fig. 3. SRRF analysis of Par-island meshwork**

**a.** Left. the average of 200 confocal images of Par3-mKate2 distribution in a cell that expresses Par3-mKate2 and Par6-GFP. Those 200 images were used for SRRF analysis (Fig. 4**a)**. The middle Panel shows the SRRF-processed image of Fig. 4**a**, which was processed with edge detection (see Methods). By this process, the continuous contours become clearly visible. Edges in the image were visualized in green. The right panel shows tracing of continuous contours of the SRRF-processed image (Fig. 4**a)** after edge detection (light blue lines). The distribution of the lengths of the continuous lines is shown as a histogram in Fig. 4**c**. Scale bar, 5 μm.

**b.** The list of means and s.d., of 7 Gaussian curve (Fig. 4**d**), combinations that best fit the density plot of the continuous contour lengths distribution shown in Fig. 4**d** (see Methods).

#### Supplementary Fig. 4

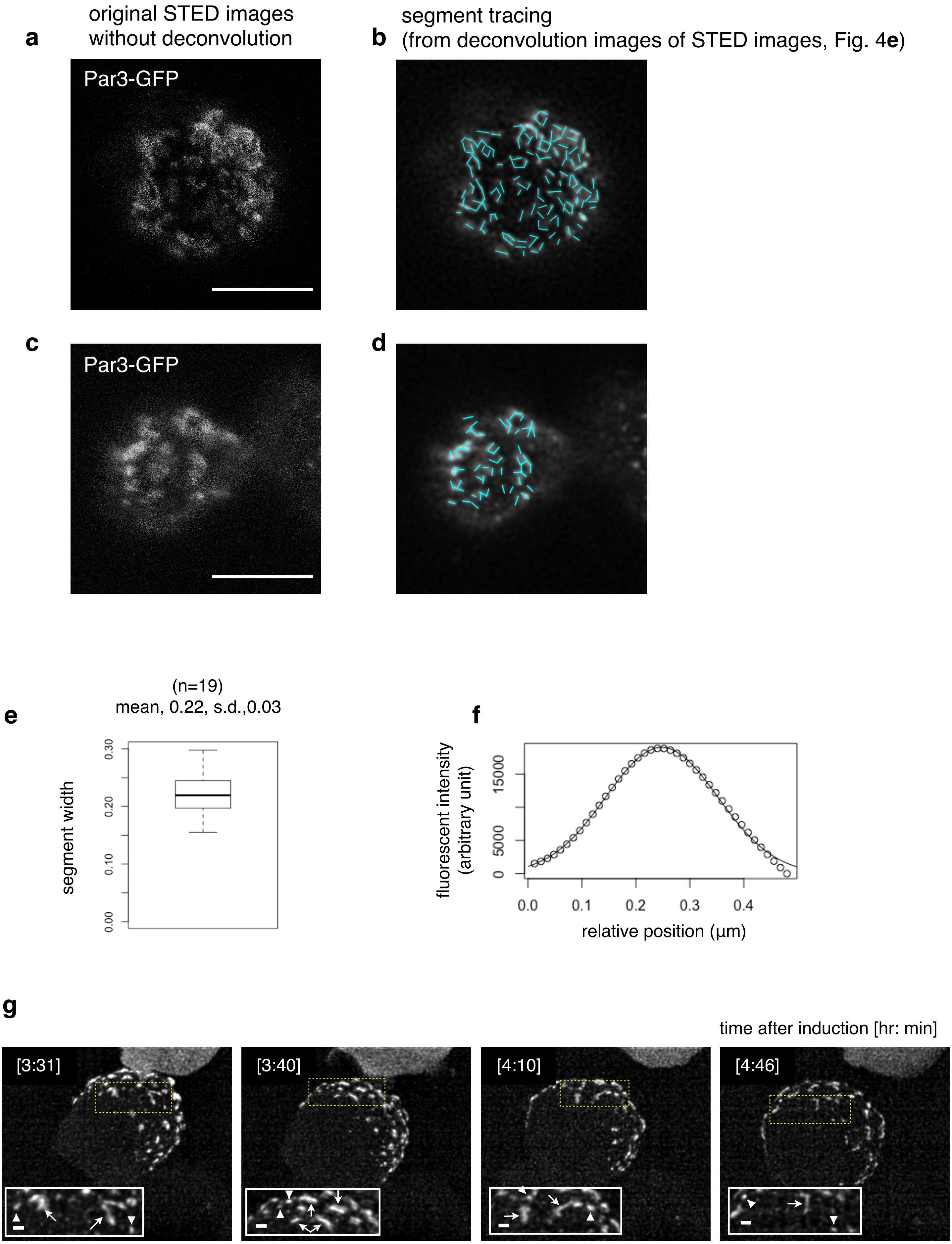

**Supplementary Fig. 4 STED analysis of Par-island meshwork**

**a.** STED image of a cell that expresses Par3-GFP. The distribution of GFP was detected via indirect immunofluorescence staining. The image on the left in Fig. 4**e** shows the deconvolved image (see Methods). Scale bar, 10 μm for (**a**-**d**).

**b.** Individual segments composing Par-islands were traced in the image (**a**) after deconvolution (Fig. 4**e**). Tracing of the segments is indicated as light blue straight lines.

**c.** STED image of a cell expressing Par3-GFP. The deconvolution of the original image is shown on the right in Fig. 4**e**.

**d.** Tracing of individual segments of Par-islands in the right-side image of Fig. 4**e**. The lengths of segmental rods detected in these 2 images (**b**,**d**) are plotted in Fig. 4**h**.

**e.** Distribution of half widths of segments composing Par-islands that were visualized by GFP staining in the two cells shown in Fig. 4a. The mean is 0.22±0.03 μm (n=19).

**f.** An example of Gaussian fitting of the fluorescence intensity distribution along the segment width visualized via immunofluorescence-staining for GFP (half width=0.24 μm).

**g.** Rod and string structures of the Par3-mKate2 in 3D time-lapse images of a cell expressing Par3-mKate2 during the period of Par-dot formation and development. Four time points were selected from Supplementary movie 2. Scale bar, 2 μm. Insets display the magnification of a part of the image. Arrowheads indicate Par-dots, arrows, rods, and strings. Scale bar, 0.5 μm.

### Supplementary Fig. 5

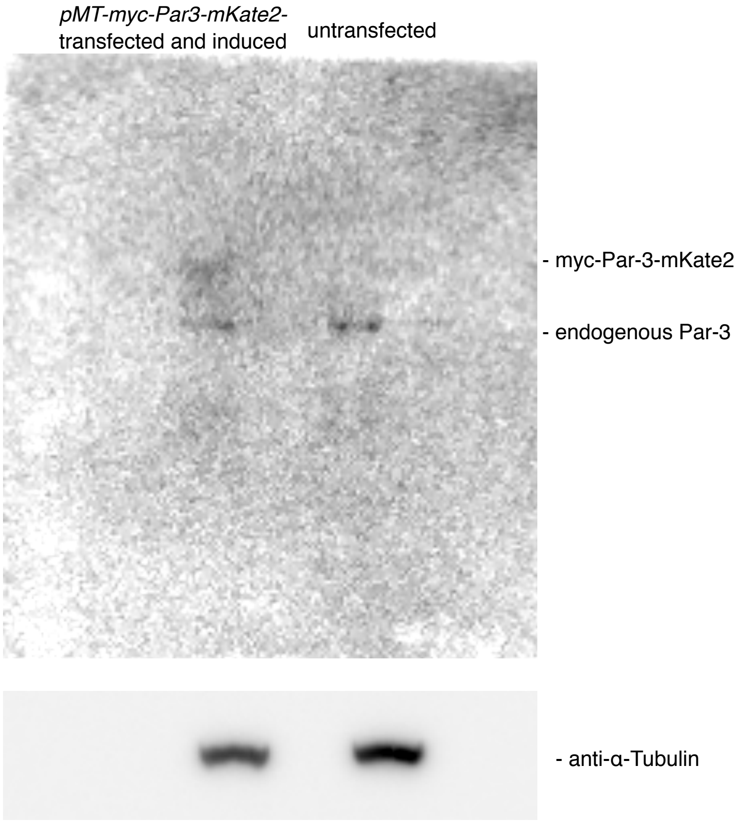

**Supplementary Fig. 5. Comparison of Par3 expression level**

Comparison of expression levels between Par3 induced by the *MT* promoter and endogenous Par3 determined via Western blotting. S2 cells were transfected with *pMT-myc-Par3-mKate2*, and induced for 8 h using $CuSO_4$ from two days post-transfection for Western blotting. Left lane: 8 h-induction of myc-Par3-mKate2 by the *MT* promoter, right lane: not transfected with *pMT-myc-Par3-mKate2*.

**List of movies**

Movie 1: 3D time-lapse movie of a polarized S2 cell monitored by Par6-GFP two days after transfection of pAct-Gal4 > UAS-myc-Par3-mKate2, pAct-Par6-GFP, and pAct-aPKC. Par-islands are clustered with dynamic movements.

Movie 2: 3D time-lapse movie of a S2 cell monitored by Par6-GFP following induction of myc-Par3-mKate2 from *Metallothionein* promoter. Induction started at time 0 by the addition of CuSO4 two days after transfection of pMT-myc-Par3-mKate2, pAct-Par6-GFP, and pAct-aPKC.

Movie 3: 3D time-lapse movie of a polarized S2 cell monitored by Par6-GFP at 8 h induction of myc-Par3-mKate2 with the co-expression of pAct-Par6-GFP and pAct-aPKC.

Movie 4: 3D movie of a polarized S2 cell stained for myc-Par3-mKate2 and Lgl at 8 h induction of myc-Par3-mKate2 with the co-expression of pAct-Par6-GFP and pAct-aPKC.

Movie 5: 3D movie of a nonpolarized S2 cell stained for myc-Par3-mKate2 and Lgl at 8 h induction of myc-Par3-mKate2 with the co-expression of pAct-Par6-GFP and pAct-aPKC. A part of the adjacent cell is included in the movie.

Movie 6: 3D time-lapse movie of a S2 cell monitored by Par6-GFP, 3 min after the addition of Latrunculin B, at 8 h induction of myc-Par3-mKate2, following two days transfection of pMT-myc-Par3-mKate2, pAct-Par6-GFP, and pAct-aPKC.
